# Receptor-linked environment-sensitive probe monitors the local membrane environment surrounding the insulin receptor

**DOI:** 10.1101/2020.12.23.424145

**Authors:** Miwa Umebayashi, Satoko Takemoto, Luc Reymond, Mayya Sundukova, Ruud Hovius, Annalisa Bucci, Paul A. Heppenstall, Hideo Yokota, Kai Johnsson, Howard Riezman

## Abstract

Functional membrane proteins in the plasma membrane are suggested to have specific membrane environments that play important roles to maintain and regulate the function of proteins. However, the local membrane environments of membrane proteins remain largely unexplored due to the lack of techniques allowing to monitor them in living cells. We have developed a method to probe the local membrane environment surrounding a membrane protein in the plasma membrane by covalently tethering a solvatochromic, environment-sensitive dye, Nile red, to a membrane protein via a flexible linker. Our direct imaging reported on the spatio-temporal properties of membrane fluidity of the local environment surrounding the insulin receptor. The local environment was distinct from the average plasma membrane fluidity and was quite dynamic and heterogeneous. Upon addition of insulin, the local membrane environment surrounding the receptor increased in fluidity in an insulin receptor-kinase dependent manner. This new technology should allow researchers to examine changes in membrane properties caused by receptor activation and devise ways to address the role of these changes in physiological processes.

## Introduction

Cellular membranes contain diverse lipid species with distinct structural and physical properties that are heterogeneously distributed in the cellular membrane, forming dynamic subcompartments with certain subsets of lipids and membrane proteins. The function and structure of integral membrane proteins are affected by the physical properties of the local membrane, composition, thickness and packing of membrane lipids, in which the membrane proteins are exposed. It has been suggested that the activities of several types of ligand-gated ion channels [1, 2], P-type adenosine triphosphatases [3, 4] and aquaporin [5] are altered by cholesterol. Cholesterol regulates the thickness, packing and fluidity of membranes as do the degree of saturation and length of lipid acyl chains [6], so lipid effects on protein function could come through direct protein lipid interactions or be exerted through biophysical properties.

Membrane lipids provide structural support to maintain the function of membrane proteins, but they also play critical roles in signal transduction. Upon stimulation, certain receptors trigger local changes in lipid composition, for example through phosphatidylinositol lipid kinases[7], which recruit downstream signalling molecules that are currently being investigated as potential therapeutic targets[8]. However, very little is known about how these pathways, along with the conformational changes of the receptors themselves, affect the local environment of the receptor and potentially its activity. Advanced imaging techniques have been able to demonstrate that there are clusters of specific proteins and lipids in membranes showing that membranes are not completely homogenous laterally [9, 10]. For example, single-molecule imaging, and fluorescence correlation spectroscopy (FCS) combined with stimulated emission depletion (STED) super-resolution microscopy evaluated the lateral diffusion of molecules of clusters and suggest the existence of nanoscale assemblies of glycosylphosphatidylinositol (GPI)-anchored proteins, cholesterol and sphingolipids [11–14]. Homo-Förster resonance energy transfer (FRET) also revealed the cholesterol-dependent clustering of GPI-anchored proteins [15, 16]. However, these fluorescence-based tehcniques reveal the dynamics of each component of lipid-protein clusters separately and it is impossible to know if they are truly in the same domains. In order to examine the local membrane environment surrounding a specific membrane protein we need other techniques.

For visualization of the membrane environment, environment sensitive dyes are widely utilized in model membranes and *in vivo*. Environment sensitive dyes sense water penetration in the membrane, exhibit a red shift in the emission spectrum depending on the local environment polarity, and membrane order is quantified by the ratio of the fluorescent intensity in two fluorescent emission regions as the generalized polarization (GP) value [17–19]. Commonly used environment sensitive dyes such as Laurdan, Di-4ANEPPDHQ and NR12S, a Nile red derivative, have hydrophobic chains and randomly reside within the membrane [20, 21].

Previously, we directly linked an environment sensitive dye, Nile red to target molecules using protein-labeling tags, the SNAP-tag [22] and the acyl carrier protein (ACP)-tag [23] to control the distribution of the environment-sensitive dye in cells, and showed that probe fluorescence depended upon attachment to the receptor [24] and we measured membrane potential in cells and in neurons [25].

Here, we directly visualized the local membrane environment surrounding the insulin receptor in living cells upon insulin stimulation by covalently linking Nile red to the receptor via a flexible tether and the ACP-tag. With proper tether length Nile red only resides in the limited area surrounding the insulin receptor by our labeling technique and the photo-stability of Nile red allows a long time-lapse observation. Thus, we can garner information about the local membrane environment of the insulin receptor with high spatial- and temporal-resolution using our system. We show that the membrane fluidity of the local membrane environment surrounding the insulin receptor is increased by insulin. The membrane fluidity change invoked by insulin requires the intrinsic tyrosine kinase activity of the insulin receptor. Meanwhile, changes in the membrane environment of the entire plasma membrane due to insulin treatment is negligible. Our findings provide evidence that the local membrane environment of the insulin receptor behaves differently from the bulk plasma membrane.

## Results

### Development of a method for probing the local membrane environment surrounding the insulin receptor in the plasma membrane

We achieved specific labelling of Nile red carrying a flexible-PEG linker with the genetically encoded ACP-tagged insulin receptor to monitor the local membrane environment of the insulin receptor. Nile red (NR) is confined to the region surrounding the receptor by direct linkage to the receptor via PEG-linker (Fig.1a). ACP-tag is a protein-labeling tag which reacts specifically with the coenzyme A (CoA) in the presence of magnesium, ACP synthase or 4’-phosphopantetheinyl transferase (SFP) synthase. To covalently label the extracellular ACP-tagged insulin receptor, we first synthesized CoA-PEG11-NR. As the membrane localization of Nile red can be determined by the distance from the ACP-tag on the receptor to the plasma membrane and the PEG-linker length of CoA-PEG-NR (Fig. 1 b), we constructed three insulin receptors with an ACP-tag at distinct positions in the α-subunit, and the ACP was inserted between Glu^664^ and Asp^665^, Gly^677^ and Leu^678^, or Lys^730^ and Thr^731^ of the receptor in constructs of 1992-ACP-IR, 2031-ACP-IR or PreCT-ACP-IR, respectively (Fig.1 c). Due to the inverted V-shaped structure of the ectodomain of the insulin receptor [26], we predicted that PreCT-ACP-IR would be the furthest ACP-tag from the plasma membrane among the three constructs even though the position of ACP-tag is inside the α C-terminal domain near the transmembrane domain (Fig.1c).

**Figure 1.**
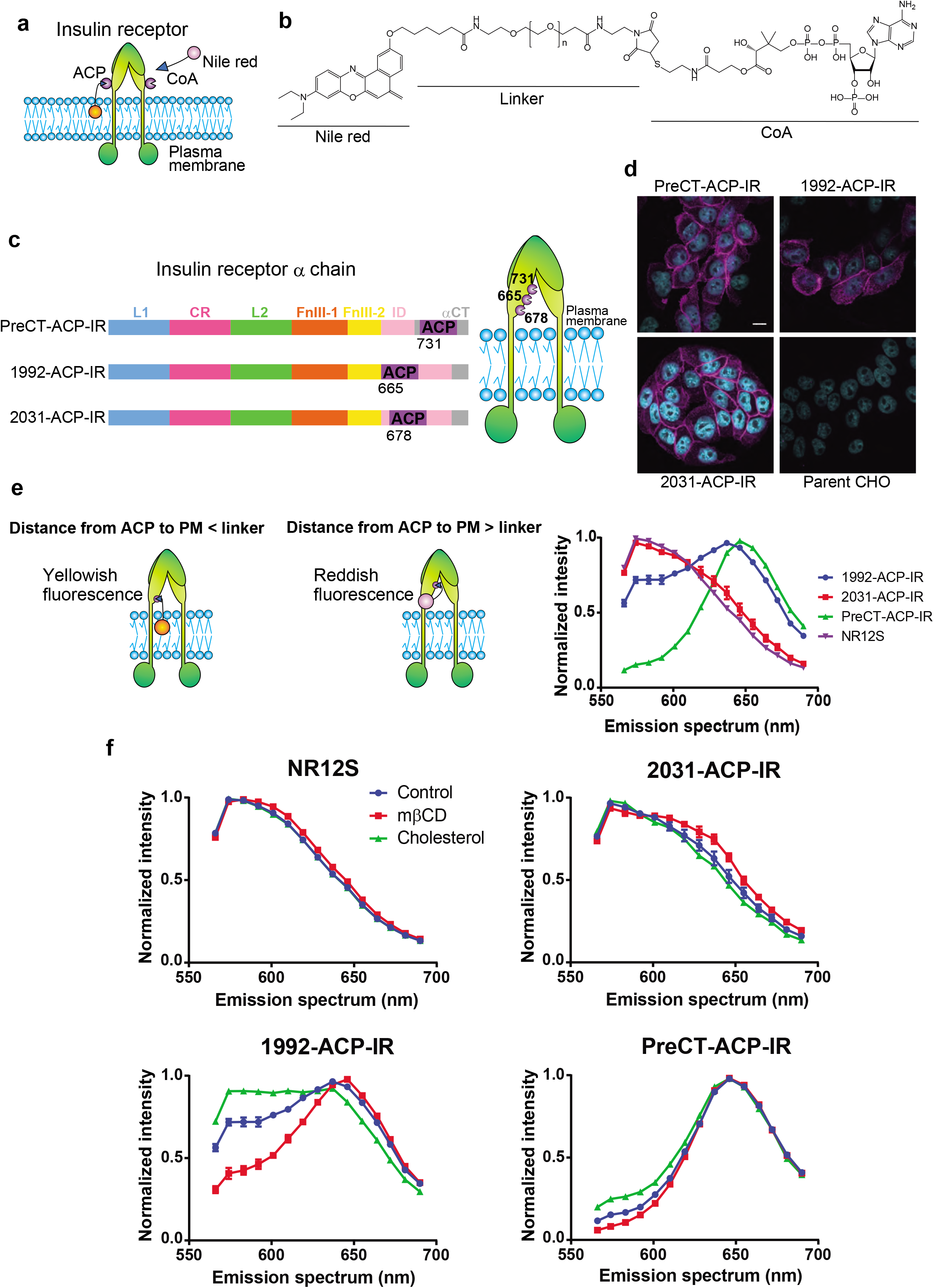
Strategy for monitoring the local membrane environment surrounding the insulin receptor and validation of membrane localization of the dye. a) Schematic image of monitoring the local membrane environment surrounding the insulin receptor using ACP-tag and CoA-derivatized Nile red. The probe is quenched in medium and Nile red is localized in the plasma membrane upon reaction with ACP-tag on the receptor, then Nile red becomes fluorescent (orange). b) Structure of CoA-PEG-Nile red with various PEG-lengths, n=5, 11, 14 and 27. c) Schematic of constructs used. Domain structures of insulin receptor α chain and the cartoon of whole insulin receptor depicting ACP-tag positions. d) Specific labelling of ACP-insulin receptor in the plasma membrane with CoA-PEG11-NR. Cyan is DAPI staining in nucleus and magenta is CoA-PEG11-NR staining. Scale bar, 10 μm. e) Emission spectral shift depending on distance from ACP-tag on the receptor to the plasma membrane. Cells expressing each ACP-IR were stained with CoA-PEG11-NR or NR12S and emission spectrum between 561 and 695 nm was measured from 5 images for 1992-ACP-IR (n=59 regions of interest (rois)), 2031-ACP-IR (n=37) and PreCT-ACP-IR (n=69), and from 7 images for NR12S (n=42). Normalized fluorescent intensities were plotted. f) Sensitivity of emission spectrum to cellular-cholesterol modification. Cell expressing each ACP-IR were treated with 10 mM of cyclodextrin for 15 min or 2.5 mM of cholesterol-cyclodextrin for 1 hr at 37 °C followed by labelling with CoA-PEG11-NR or NR12S, then the emission spectrum was measured. Data represent the mean ± S.E.M from 5 images (n=57-59 rois for 1992-ACP-IR, n=36-37 for 2031-ACP-IR and n=69-75 for PreCT-ACP-IR, and the mean ± SE from 28 images of NR12S (n=168 rois for each sample).

We first confirmed that the plasma membrane staining with CoA-PEG-Nile NR was dependent upon expression of the ACP-tagged IR. The plasma membrane staining with CoA-PEG11-NR was observed in ACP-IR expressing cells, whereas there was no signal of CoA-PEG11-NR detected in parent cells (Fig. 1d).

It is essential that the Nile red moiety is inserted into the plasma membrane upon binding to the ACP-IR in order to monitor the local membrane environment of the receptor. We next validated the membrane localization of Nile red of phosphopantatheine(PP)-PEG11-NR linked to each ACP-IR by fluorescence spectroscopy. Nile red is a solvatochromic dye which exhibits a red shift depending on water presence in the environment and we expected that Nile red could reach to the plasma membrane and emit yellowish fluorescence when the distance between ACP-tag on the receptor and the plasma membrane is shorter than the linker length of CoA-PEG11-NR, whereas Nile red of PP-PEG11-NR linked to ACP-insulin receptor whose ACP-tag is far from the plasma membrane would interact somewhere on the receptor and emit reddish fluorescence. These prediction were confirmed by our measurements (Fig. 1e).

The emission maximum (λ_max_) of PP-PEG11-NR linked to the site closest to the membrane, 2031-ACP-IR, was 574 nm which was comparable to that of NR12S, a Nile red derivative with an amphiphilic anchor exclusively bound to the outer leaflet of the plasma membrane [20]. On the other hand, PP-PEG11-NR attached to the middle site, 1992-ACP-IR, or the furthest site, PreCT-ACP-IR, showed peaks at 637 or 646 nm, respectively (Fig. 1e), and these are higher than the emission peak of Nile red in liquid disordered membranes, around 620 nm [20]. This strongly suggests that Nile red of PP-PEG11-NR linked to 1992-ACP-or PreCT-ACP-IR binds somewhere on the receptor rather than being in the membrane.

To provide further evidence for the membrane localization of PP-PEG11-NR connected to ACP-IR, the sensitivity of the emission spectrum to cellular-cholesterol modification was tested. The emission peak at 574 nm was decreased and a shoulder around 620 nm was increased by cholesterol-depletion with cyclodextrin. Conversely, the peak at 574 nm was slightly increased and the shoulder peak around 620 nm was lowered by cholesterol-loading when PP-PEG11-NR was linked to the closest 2031-ACP-IR. Similar, but less pronounced patterns were observed with NR12S. On the other hand, there was no emission spectral shift by cellular-cholesterol modification with PP-PEG11-NR attached to the furthest PreCT-ACP-IR (Fig. 1f). These results indicate that Nile red of PP-PEG11-NR responds to cholesterol concentration and therefore is in the plasma membrane when the dye is linked to 2031-ACP-IR. Interestingly, the largest changes were obtained with the 1992-ACP-IR construct. Increasing membrane cholesterol seemed to increase its insertion into the membrane with a corresponding increase in the peak at 574 nm. Increasing cholesterol may affect lipid packing and membrane thickness, membrane properties which may increase the insertion or stability of insertion of the dye into the membrane under these circumstances where the tether brings the Nile Red chromophore close to the membrane.

The experiment above suggests that some portion of the NR is dynamically close enough to the membrane to insert into the bilayer when it is tethered to the 1992-ACP-IR. In order to test this, we examined the effect of PEG-linker length, using CoA-PEG5, 14, 27-NR. The λ_max_ of all of PP-PEG-NR constructs attached to 2031-ACP-IR was 574 nm, consistent with the membrane localization of Nile red regardless of the PEG-linker length (Fig. 2a). On the other hand, Nile red of PP-PEG-NR linked to the 1992-ACP-IR exhibited large emission shifts according to the PEG-linker length. The peak of PP-PEG5-NR was 646 nm and the peak of PP-PEG27-NR was 574 nm similar to NR12S, suggesting that Nile red with the shortest PEG5-linker binds the receptor, while Nile red with the longest PEG27-linker sits in the membrane (Fig.2 b). One can see that membrane insertion probably begins with a linker length of 11, but improves as the linker is extended, consistent with the previous explanation concerning membrane properties.

**Figure 2.**
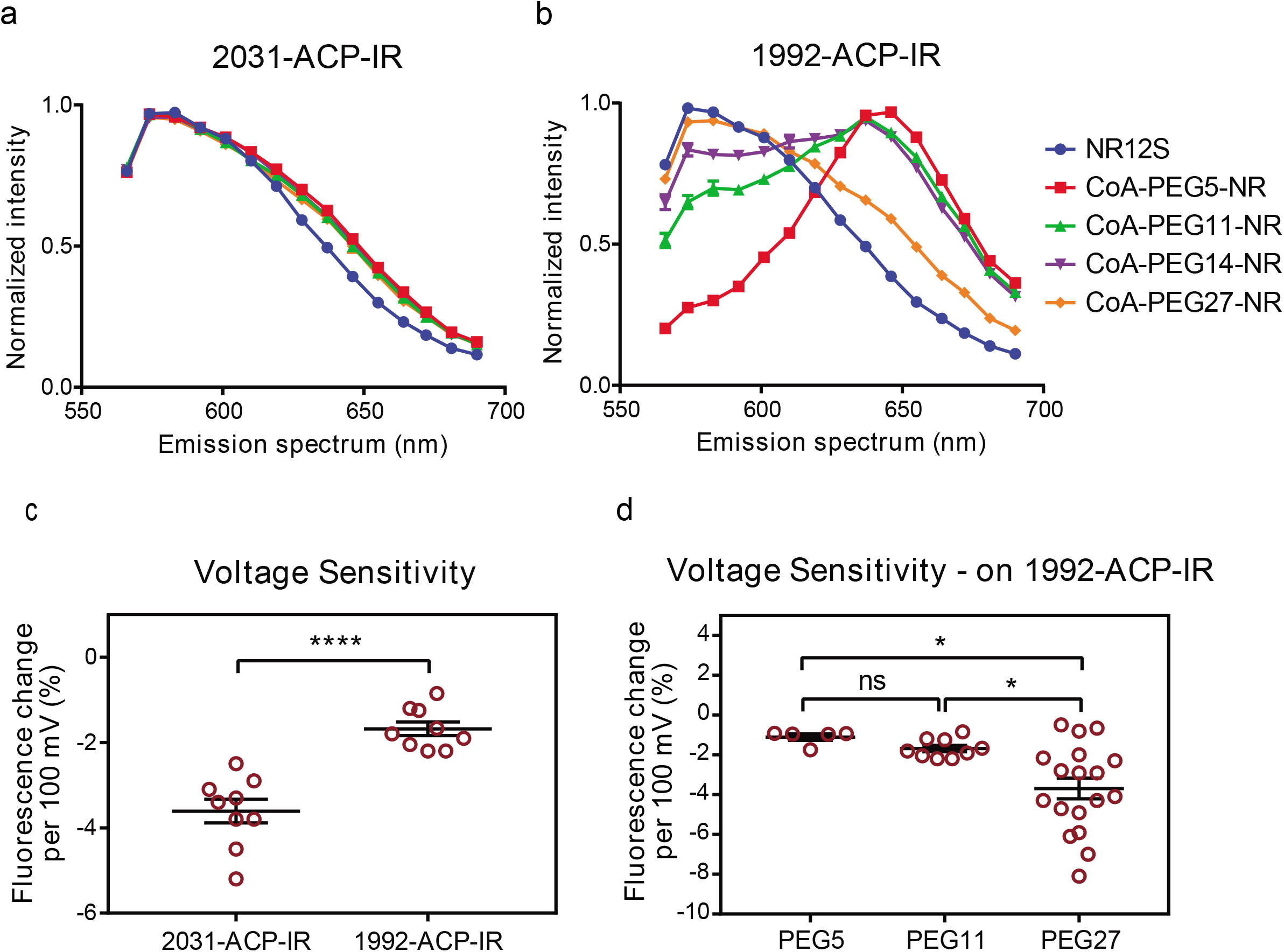
Effect of emission spectrum of CoA-PEG-NR bound ACP-IR to PEG-linker length and sensitivity test to membrane-voltage change. a and b) Sensitivity of emission spectrum to PEG-linker length. Cells expressing 2031-ACP-IR (a) or 1992-ACP-IR (b) were labelled with CoA-PEG_n_-NR (n=5, 11, 14 or 27) or NR12S and emission spectrum was analysed (each sample: n=75 from 5 images). c) Membrane depolarization led to decreased fluorescence intensity of PP-PEG11-NR bound 2031-ACP-IR.2031-ACP-IR (n=9) or 1992-ACP-IR (n=9) expressing cells labelled by CoA-PEG11-NR and the change in fluorescence intensity induced by membrane depolarization (100 mV) using patch-clump was recorded. Data are means ± S.E.M. d) Correlation between the membrane localization of dye and voltage sensitivity. Cells expressing 1992-ACP-IR were stained with CoA-PEG5-, 11-or 27-Nile red and the fluorescent intensity was measured upon membrane depolarization (100 mV). Data represent means ±S.E.M. of n=5 (PEG5), 9 (PEG11) and 18 (PEG27).

Recently, we have demonstrated that NR12S or PP-PEG5- and 11-NR linked to ACP-tag anchored to the plasma membrane via a GPI were sensitive to membrane voltage change using a patch-clamp technique in HEK cells and neurons [25]. Using this technique, we further confirmed the membrane insertion of Nile red of PP-PEG11-NR linked to ACP-IR by the voltage-sensitivity test. The fluorescent intensity of PP-PEG11-NR attached to 2031-ACP-IR was decreased – 3.4 ± 0.3 % (n= 9) when the membrane voltage was increased to 100 mV, however PP-PEG11-NR bound 1992-ACP-IR showed no significant difference (Fig. 2c). We also tested voltage sensitivity of PP-Nile red with various length PEG-linkers in cells expressing 1992-ACP-IR. The fluorescent intensity per 100 mV was decreased with the length of PEG-linker and the delta % changes were −0.96 ± 0.16 for PEG5 (n=5), −1.8 ± 0.2 for PEG11 (n=9) and −3.5 ± 0.5 for PEG27 (n=18) (Fig. 2d). Taken together with the results using fluorescence spectroscopy, there is a correlation between the membrane localization of the dye and voltage sensitivity, showing that Nile red of PP-PEG27-NR attached to 1992-ACP-IR is indeed embedded in the membrane, whereas PP-PEG5 interacts the receptor itself. Consistent with the results above, Nile red tethered with 11 PEG may partially insert into membrane.

#### The local membrane environment surrounding the insulin receptor becomes more disordered upon insulin stimulation

The experiments above confirmed that Nile red moiety of PP-PEG-NR resides in the plasma membrane upon binding to 2031-ACP-IR by fluorescent spectroscopy and voltage sensitivity tests. Next, we checked the activity of 2031-ACP-IR upon insulin treatment. The basal level of tyrosine phosphorylation of 2031-ACP-IR without insulin addition was similar in the presence or absence of CoA-PEG11-NR. Upon insulin addition, tyrosine phosphorylation was increased on the 2031-ACP-IR, in both unlabeled and PP-PEG11-NR-labeled forms, showing that the modified receptor is still active and that the attachment of the PP-PEG-NR with this long linker did not significantly affect the tyrosine kinase activity of 2031-ACP-IR (Fig. 3a).

**Figure 3.**
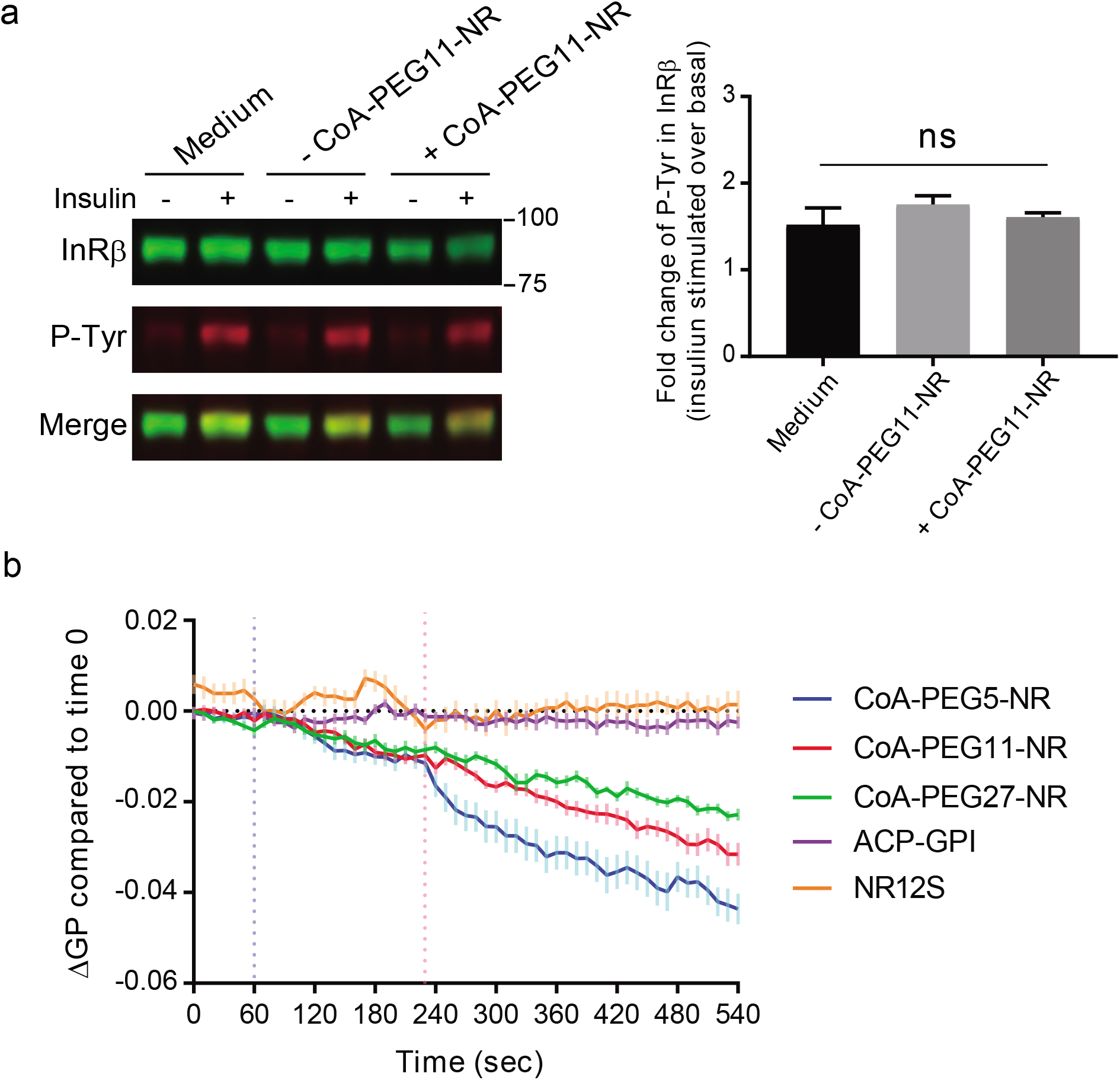
PP-PEG5-NR linked ACP-insulin receptor detects the change in the local membrane environment surrounding the insulin receptor caused by insulin. a) Tyrosine-phosphorylation of 2031-ACP-IR upon insulin in the presence or absence of attachment of PEG11-NR. 2031-ACP-IR expressing cells were serum-starved for 2 hr followed by incubation with serum-free medium, medium including SFP synthase, or medium including SFP synthase and CoA-PEG11-NR for 30 min at room temperatures, then cells were stimulated by 100 nM of insulin at 37°C for 5 min and phosphorylation of 2031-ACP-IR was accessed by western blotting analysis using α-insulin receptor β antibody (left upper bands) and α-phospho-tyrosine antibody (left middle bands). Tyrosine-phosphorylation levels are presented as the mean fold increase over the basal ±S.E., n=7-8, for each condition. Statistical significance was calculated by one-way ANOVA, ns, not significant. b) Time course of mean delta GP change compared to time 0. After 2 hr serum starvation, cell stably expressing 2031-ACP-IR were labelled with CoA-PEG5-, 11-, or 27-NR or NR12S and cells transiently expressing ACP-GPI were stained with CoA-PEG11-NR. Spectral imaging was performed every 10 sec and the control solution and insulin solution were added at 60 and 240 sec, respectively. GP value was calculated with two emission regions, 561-597 nm and 605-641 nm, and the delta GP compared to the time 0 at each time point was plotted. Data represent the mean ± S.E.M of from 19 time-lapse image sets for CoA-PEG5-NR labeled ACP-IR, 22 for PP-PEG11-NR labeled ACP-IR, 17 for PP-PEG27-NR labeled-ACP-IR, 12 for labeled with NR12S, and 19 for PP-PEG11-NR labeled ACP-GPI.

Next, we observed the local membrane environment of the insulin receptor upon insulin signaling using 2031-ACP-IR labelled with PP-PEG5-, 11-, or 27-NR. We also observed the local membrane environment of ACP-GPI linked to PP-PEG11-NR and the overall properties of the plasma membrane with NR12S. A GP value was calculated to assess the relative membrane order and the delta GP compared to time 0 was plotted at each time point (see methods). No major changes in membrane order were detected using NR12S upon insulin addition, indicating that the average properties of the plasma membrane did not change upon insulin binding and signaling. Likewise, the local membrane environment surrounding ACP-GPI was almost unchanged upon on insulin treatment as the GP value was constant through the whole observation. Strikingly, the GP value clearly dropped upon insulin treatment when the 2031-ACP-IR was labeled with PP-PEG5-NR. This shows that the membrane environment immediately surrounding the insulin receptor became more disordered upon insulin binding and signaling, distinct from the bulk membrane. As the length of the PEG linker was increased situating the NR further away from the receptor, the impact of insulin treatment on the GP value was weaker, but still remained even with PEG27 (Fig. 3b). These results suggest that the changes invoked by insulin treatment happened in the immediate vicinity of the insulin receptor and that the further away the NR was positioned the less apparent the effects. These results further support that our measurements are relevant to the immediate membrane environment of the receptor and suggest that by increasing linker length that we can determine distance effects.

The actual GP change induced by insulin was small and therefore, it was difficult to explore which area of the local membrane environment surrounding the insulin receptor shows significant change after insulin treatment in pseudocolor GP images (Fig. 4a, supplementary fig. 2a and supplementary video 1). We thus employed zero-mean cross correlation (ZNCC), a similarity measure metric often used for template-matching, to visualize statistical GP changes between the time-lapse image and the averaged image of control-treatment period from 60 sec to 230 sec. The range of ZNCC value is −1 to 1 and 1 means the perfect matching to the reference image. To take into consideration cell migration and membrane movement through the time-lapse observation, a local region was set to 19×19 pixels in each frame image so that the impact of cellular movement can be canceled inside the local region for calculating ZNCC (methods, ZNCC). We could clearly see changes in the receptor membrane environment after insulin treatment in ZNCC images. Interestingly, the cellular responses to insulin treatment were quite heterogeneous. There were cells that responded strongly and rapidly to insulin, while some cells showed delayed reactions and a few cells barely responded to insulin (Fig. 4 b,d, supplementary fig. 2b, d and supplementary video 2). We also noticed that the environment surrounding receptors in certain parts of the plasma membrane changed more upon insulin stimulation. In order to identify receptor environments that responded more or less intensely to insulin the top 1000 pixels showing biggest ZNCC changes or bottom 1000 pixels showing smallest ZNCC changes were analysed in all images. All cells were moving through the observation and some cells showed extensive membrane blebbing and the top 1000 pixels often included regions of extensive membrane blebbing (Fig. 4c, supplementary fig. 2 c, e, and supplementary video 3). Kymographs were made for regions containing the top (Fig.4 d 1, 2, 4, 5 and 6) or bottom 1000 pixels (Fig. 4d 3). The kymographs of 1, 2, 4 and 6 include regions that had protrusions (Fig. 4b,supplementary fig. 2d and supplementary video 2). ZNCC kymographs clearly showed that changes were triggered by insulin treatment and the local membrane environment of the insulin receptor was quite heterogeneous. Some areas exhibited significant change just after insulin stimulation (Fig. 4 d1, 2, 4 and 6), whereas changes were observed late in the position 5 and the position 3 exhibited negligible change upon insulin stimulation (Fig. 4d). The whole time-lapse image set of GP, ZNCC or top/bottom 1000 is shown in the supplementary figure 2a, b, c and supplementary video 1, 2 or 3, respectively.

**Figure 4.**
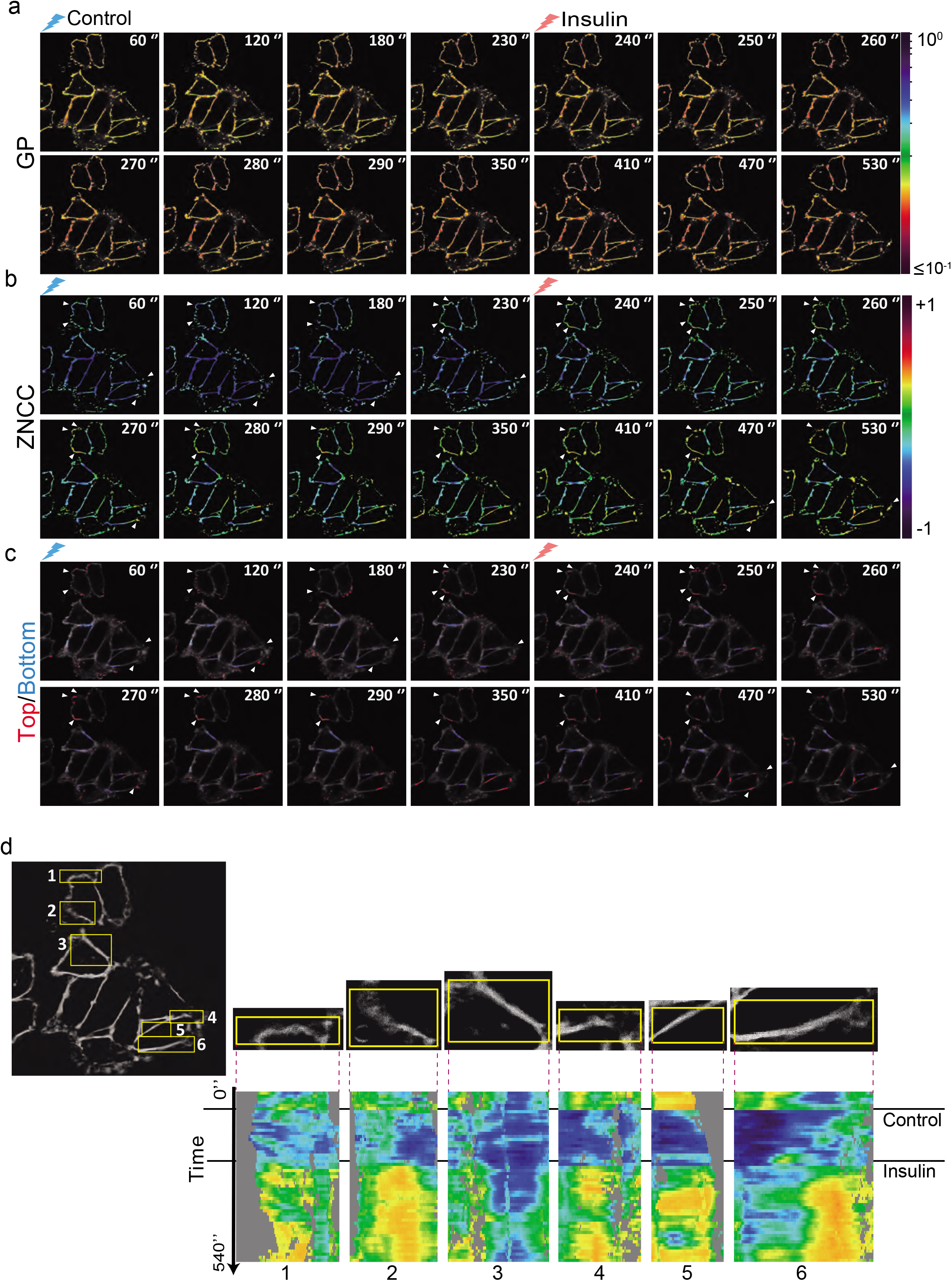
ZNCC images reveal that the local membrane environment of the insulin receptor is quite dynamic and heterogeneously react to insulin. A representative time-lapse data set of pseudocolor GP images filtering the median filter (3 × 3 pixels) (a), pseudocolor ZNCC images (b), and top/bottom 1000 pixel images exhibiting biggest 1000 pixels of ZNCC changes (red) and smallest 1000 pixels of ZNCC changes (blue) (c). a) The pseudocolor GP images are depicted by logarithmic scale from 10^-1^ to 10^0^. The GP value less than 10^-1^ is displayed in the same color as 10^-1^. b,c) Arrows are included to facilitate following specific features through the time-course. This time-lapse image set was one data set utilized for delta GP change compared to time 0 in Fig. 3b. d) Kymographs of chosen areas show time-course ZNCC changes. The horizontal dimension represents the averaged ZNCC inside the rectangular area, and the vertical dimension represents time (from t = 1 to t = 55).

#### Tyrosine kinase activity is required for GP change in the local membrane environment surrounding the insulin receptor by insulin treatment

We showed the GP change caused by insulin stimulation occurred only in the vicinity of the insulin receptor, whereas neither the GP in the entire plasma membrane nor the local membrane environment of ACP-GPI was altered. These results suggest that the GP change in the immediate membrane environment surrounding the insulin receptor reflects local events catalyzed by the intrinsic tyrosine kinase activity of the receptor. To confirm this possibility, we constructed the kinase-dead mutant of 2031-ACP-IR with replacement of Lys 1045 in the ATP binding site of the β subunit by Arg [27, 28]. Tyrosine-phosphorylation of the mutant insulin receptor was completely blocked upon insulin stimulation (Fig. 5a). We compared the delta GP change between the wild-type 2031-ACP-IR and kinase-dead 2031-ACP-IR labeled with PP-PEG5-NR. In this experiment, differential delta GP with insulin-treatment of each sample, wild-type (WT) and kinase-dead (KD) mutant insulin receptor, against the basal delta GP of each sample with control-treatment set to 0 is plotted. Before the insulin treatment at 120 sec, the delta GP of both WT and KD insulin receptor remained at the basal level, whereas it went down immediately after the insulin stimulation in the WT but the KD stayed at pre-insulin levels (Fig.5b).

**Figure 5.**
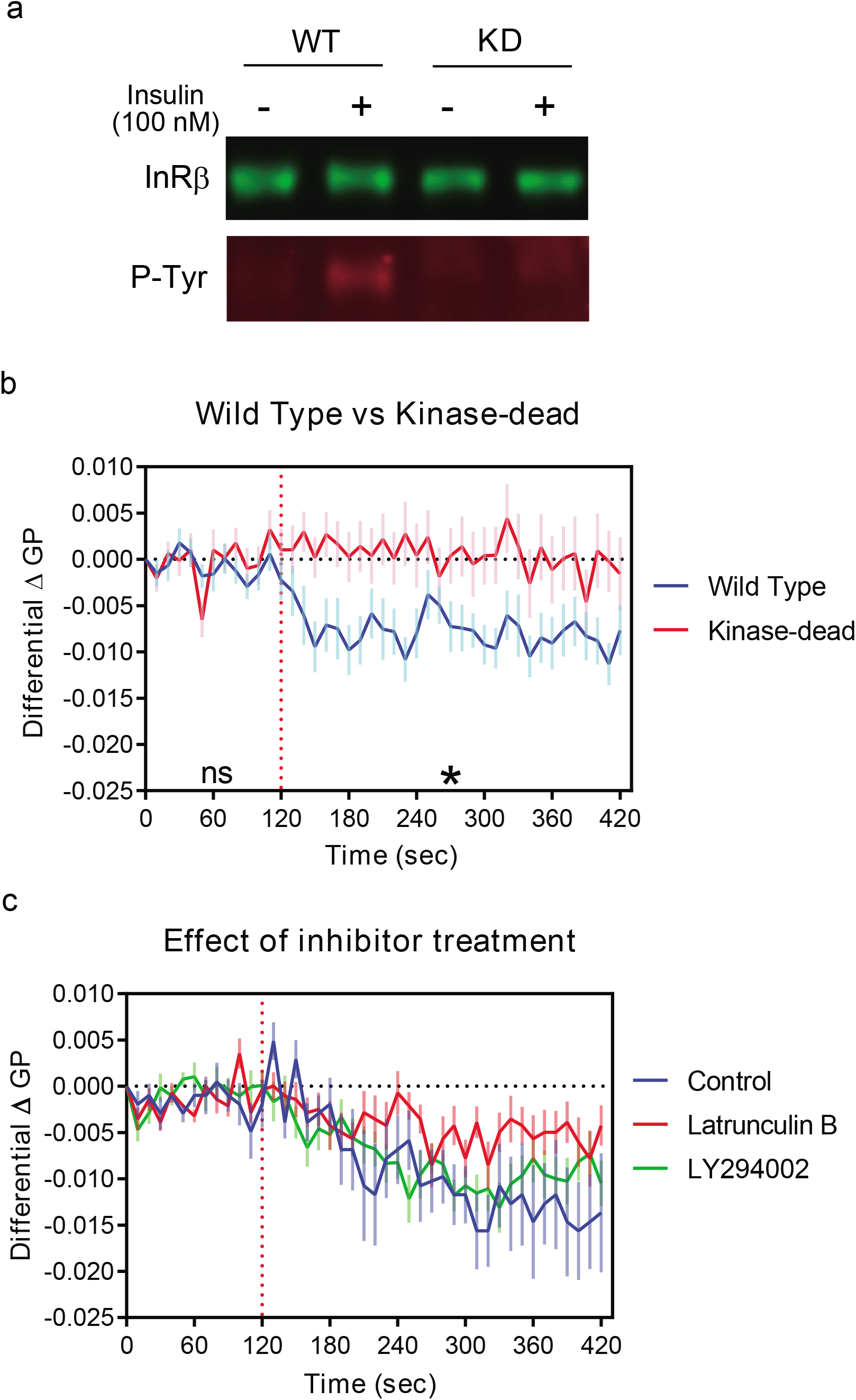
GP changes in the local membrane environment surrounding the insulin receptor invoked by insulin require the tyrosine kinase activity of the insulin receptor. a) Tyrosine-phosphorylation of the insulin receptor was inhibited upon insulin treatment in the kinase-dead mutant. Cells stably expressing the WT 2031-ACP-IR (left) or the KD mutant 2031-ACP-IR (right) were serum-starved for 2 hr, then treated with 100 nM of insulin at 37 °C for 5 min. Tyrosine-phosphorylation of ACP-IR was analyzed by western blotting using α-insulin receptor β antibody (upper bands) and α-phospho-tyrosine antibody (lower bands). b) The differential delta GP suggested that the local membrane environment of the KD mutant insulin receptor showed no GP change upon insulin treatment (red), whereas the GP in the local membrane environment surrounding the WT insulin receptor dropped after insulin stimulation (blue). Stably expressing the WT 2031-ACP-IR or the KD 2031-ACP-IR cells were serum-starved for 2 hr, followed by labeling with CoA-PEG5-NR, then spectral imaging was performed and the control solution or insulin solution were added at 120 sec. The differential delta GP with insulin treatment of both WT and KD over the basal control-treated delta GP of each sample was plotted. The basal control-treated delta GP was estimated as the mean delta GP of 11 time-lapse data sets for both WT and KD with. Data represent means ± S.E.M. of 13 time-lapse data sets for WT and 15 for KD. Changes between WT and KD were statistically analyzed; during 0-120sec, no significance, after the insulin treatment at 120 sec, the change was significant *, p<0.05. c) The differential delta GP was analysed in cells stably expressing WT 2031-ACP-IR in the presence of dimethyl sulfoxide (DMSO) (control), Latrunculin B or LY 294002. The basal control-treated delta GP was calculated as the mean of 8 time-lapse data for control, 11 for Latrunculin B and 9 for LY294002. Data present means ± of 8 time-lapse data for control, 11 for Latrunculin B and 10 for LY294002.

Activation of the insulin receptor tyrosine kinase phosphorylates the insulin receptor and the insulin receptor substrate proteins, leading to the recruitment of phosphatidylinositol 3-kinase (PI3K) to the receptor, then phosphatidylinositol (3,4,5) triphosphate (PIP_3_) is produced [29]. Therefore, we examined the effect of PIP_3_ production on the GP change induced by insulin treatment by using LY294002, a PI3K inhibitor. The phosphorylation of Akt which is downstream of PI3K was completely blocked by LY294002 treatment (supplementary fig. 4), however, the GP change induced by insulin was similar to the untreated control (Fig. 5c). Therefore, it is unlikely that PIP_3_ production is required for these rapid changes in the local membrane environment.

The actin cytoskeleton is considered to be an important factor that can influence membrane organization [30]. To explore the involvement of actin cytoskeleton in the GP drop evoked by insulin, we perturbed the actin cytoskeleton using Latrunculin B. The disruption of filamentous actin structures by Latrunculin B (supplementary fig. 5) partially prevented the GP decrease in response to insulin (Fig. 5 c), suggesting that actin may play a role in these rapid local membrane environment changes.

### Discussion

We have established a novel method to visualize the local membrane environment surrounding the insulin receptor and have examined its changes due to insulin signalling. In our method, the insulin receptor is covalently tethered to an environment sensitive dye, Nile red, using a flexible PEG-linker and the ACP tag. The hydrophilicity of CoA and PEG-linker prevents the dye from crossing the plasma membrane and this is critical for specific labelling of the ACP-IR with PP-PEG(n)-NR in the plasma membrane[24]. Additionally, the enzymatic procedure for attachment of the fluorescent reporter to the ACP tag requires addition of an enzyme, further restricting the attachment to the plasma membrane. To monitor the local membrane environment surrounding the receptor in the plasma membrane, the Nile red was linked to ACP-IR through a flexible tether allowing the NR to reside in the plasma membrane. To fulfil this requirement, the ACP-tag on the ectodomain of the receptor needs to be located close enough to the plasma membrane that the length of

PEG-linker can be adjusted to bridge this critical distance. We provided several lines of evidence to prove that the NR was embedded in the membrane. First, the λ_max_ of Nile red in was around 574 nm corresponding to the expected values for NR in ordered membranes (580 nm) as opposed to disordered membranes (610 mm)[20] similar to that seen previously and with the membrane probe NR12S. Second, the fluorescence spectrum of the NR was sensitive to changes in amounts of cholesterol. Third, the NR fluorescence was sensitive to a change in membrane potential using the patch clamp technique. Furthermore, when the probe was predicted to not reach the membrane, with the PreCT-ACP-IR, the fluorescence spectrum peaked at 646 nm indicating that the NR was not membrane embedded and accordingly was insensitive to cholesterol depletion or addition. It is perhaps a bit surprizing that the NR fluorescence was detected using this construct as NR is normally only very weakly fluorescent in aqueous solution. Part of the reason why NR does not fluoresce well in water is that it aggregates. When aggregation is prevented with cyclodextrin the emission spectrum[31] is quite similar to what we see with the PreCT-ACP-IR-bound NR, consistent with the protein bound NR being in an aqueous environment. Our probes are probably much less subject to aggregation than NR due to the hydrophilic moieties that we have attached to them.

The experiments with the 1992-ACP-IR revealed other aspects and a potential use of our technology. At short PEG-linker lengths the NR is not embedded in the membrane, but as the linker length increases the NR inserts into the plasma membrane and fluoresces in a similar manner as when attached to the 2032-ACP-IR (Fig. 2a and b). Thus by varying the linker length we can bridge the distance between the site of attachment on the protein and the membrane, meaning that our probes can function as molecular rulers to measure the distance between sites on membrane proteins and the membrane. We also saw that with increased linker length the changes in membrane properties due to insulin signalling were reduced (Fig 3b) allowing us to measure distance effects with our probes. Interestingly, when the PP-PEG11-NR was linked to the 1992-ACP-IR we found two emission peaks, a major peak at 637 nm and a minor peak at 574 nm, indicating the majority of Nile red was bound to the receptor and a part of Nile red was in the membrane (Fig. 1e). Interestingly, the two peaks became approximately equal when cholesterol-loading of the membrane was performed. When cholesterol was depleted with cyclodextrin the membrane bound peak was virtually eliminated (Fig. 1f). These results showed that the proportion of the NR moiety of PP-PEG11-NR attached to the PreCT-ACP-IR that was inserted into the plasma membrane was increased by cholesterol-loading and decreased by cholesterol-depletion. Two explanations are possible to account for this behaviour. First, the membrane thickness could have been increased by the cholesterol content in the membrane bringing it closer to the Nile red. Second, the increased cholesterol content of the membrane could have stabilized the dynamic insertion of the Nile red into the membrane. In any event, suggest that functional distances of sites on membrane-associated proteins to the plasma membrane could be modulated by membrane properties.

Our results observed using the 2031-ACP-IR labelled by PP-PEG-NR with various PEG-linker length showed that the GP value in the local membrane environment surrounding the insulin receptor was decreased upon insulin stimulation, and the biggest impact of insulin treatment was obtained by PP-PEG5-NR which has the shortest PEG linker. Moreover, neither the entire plasma membrane stained by NR12S nor the local membrane environment surrounding the ACP-GPI labeled with PP-PEG11-NR were altered by insulin (Fig. 3b). These results suggest that the change of membrane environment caused by insulin occurs only in the vicinity of the insulin receptor. The changes in the local membrane environment induced by insulin were completely blocked in the kinase dead receptor (KD 2031-ACP-IR. Fig. 5b), demonstrating that the local membrane environment change requires the intrinsic tyrosine kinase upon insulin binding and probably depends upon signalling. The insulin receptor is a disulphide-linked (αβ)_2_ homodimer, comprised of heterodimers containing α- and β-subunit, and the insulin receptor undergoes the conformation changes by insulin binding. Detailed conformation changes of the whole insulin receptor driven by insulin have been proposed by singleparticle electron microscopy [32, 33]. The symmetric ectodomain of the insulin receptor is converted from an inverted U-shaped conformation into a T-shaped conformation upon insulin binding that brings the well-separated two transmembrane domains together, then facilitates autophosphorylation of the tyrosine kinase domains in the cytoplasmic tail. To understand the GP decrease in the local membrane environment of the insulin receptor caused by insulin we compared the emission spectrum of PP-PEG5-NR linked to the 2031-ACP-IR at time 0, at 230 sec, just before the insulin treatment and 540 sec, the last time point after insulin treatment. The emission spectra of 0 and 230 sec were identical, whereas the peak around 580 nm was decreased in the emission spectrum of 540 sec (supplementary fig.1). This indicates that the local membrane environment surrounding the insulin receptor gets more disordered upon insulin stimulation. The conformational change invoked by insulin may disturb the membrane layer and allow water penetration surrounding the insulin receptor. Certain conformational changes of the insulin receptor are independent of kinase activity, whereas others are dependent on autophosphorylation [34]. Since the kinase dead mutant could not induce the changes in membrane environment upon insulin binding, either the kinase-dependent conformational change or insulin receptor signalling is probably directly required for these changes.

In order to identify the areas that showed clear GP change in the local membrane environment surrounding the insulin receptor, we employed ZNCC. Due to the large local region 19×19 pixels, our ZNCC analysis succeed in demonstrating time-dependent statistical GP-change and ignoring morphological changes like membrane blebbing and ruffling. The plasma membrane in the position 1 and 2 (Fig. 4c, b arrows, d and video 2) showed membrane blebbing and ruffling through the whole observation, but ZNCC changes were observed after insulin pulse at 240 sec. ZNCC kymographs revealed that the response of the local membrane environment surrounding the insulin receptor to insulin was quite heterogeneous. Certain regions of the plasma membrane including membrane blebs showed clear changes upon insulin stimulation and the timing of response to the insulin varied depending on regions (Fig. 4 b,d and supplementary fig. 2d). Thus our technique might be able to resolve local insulin effects in different plasma membrane domains.

Our results using a PI3K inhibitor also demonstrate that the GP change in the local membrane environment of the receptor are not due to PIP_3_ production surrounding the receptor nor the downstream signalling activated by PI3K after insulin stimulation (Fig. 5c), but we cannot rule out that the local membrane environment change could be caused by gathering of the substrate, phosphatidyl inositol (4,5) bisphosphate, to the receptor before conversion to PIP_3_.

In contrast to PIP_3_ signaling, the actin cytoskeleton was required for the optimal insulin-driven GP change in the local membrane environment of the insulin receptor (Fig. 5c). It has been reported that the IR is compartmentalized in plasma membrane domains enriched in cytoskeletal components such as microvilli or the caveolae neck in the plasma membrane and relocates to be internalized upon insulin stimulation [28, 35, 36]. The direct interaction of IR with actin cytoskeleton was observed in 3T3-L1 adipocytes [36], whereas the indirect interaction via actin-binding protein, filamin, was suggested in HepG2 cells [37]. Therefore, IR-actin cytoskeleton interaction might be necessary for maintenance and formation of the specific membrane environment of the IR in insulin signalling. The local membrane environment of the insulin receptor could be mediated by other mechanisms involving actin filaments. The insulin receptor has been suggested to form actin-dependent clusters upon insulin stimulation [38] and actin also affects clustering of GPI-anchored proteins [15, 16]. Furthermore, membrane properties are affected by self-organizing actin patterns [30]. Actin might play multiple roles in the regulation of the local membrane environment of the insulin receptor during signalling.

Our system should be applicable to examine the local membrane environment of other membrane proteins in the plasma membrane. Indeed, our system recognizes differences in the local membrane environment between 2031-ACP-IR and WT ACP-GPI, and furthermore between WT ACP-GPI with two saturated fatty chains and mutant ACP-GPI deficient in PGAP3- and PGAP2-mediated fatty acid remodeling bearing un unsaturated fatty chain[39]. The local membrane environment of WT ACP-GPI is more ordered than that of 2031-ACP-IR and PGAP2/3 mutant ACP-GPI, whereas the averaged membrane environment of the plasma membrane examined with NR12S in cells expressing WT ACP-GPI or PGAP2/3 mutant ACP-GPI (supplementary fig. 6) was identical. These results also provide evidence that our system is a good tool to monitor the local membrane environment of a target membrane protein.

In order to observe the local membrane environment of other membrane proteins using our system, consideration should be given to the location of the ACP-tag on the protein and PEG-linker length of CoA-PEG-NR to make sure that the breach to the membrane can be overcome. Of course structural information on the protein of interest would facilitate this strategy, but our technology also has potential to obtain information of protein-membrane distances independent of prior structural information. Care should be taken to ensure that the protein constructs made are functional.

## Methods

### Materials

CoA-PEG-Nile reds were synthesized as described previously [25] and NR12S was a gift from Dr. Klymchenko [20]. SFP synthase was purchased from NEB. Methyl beta cyclodextrin was purchased from CycloLab. Cholesterol-water soluble, human reconstitute insulin, Complete EDTA-free protease inhibitor cocktails and Latrunculin B were purchased from Sigma-Aldrich. Pierce™ BCA Protein Assay Kit, NuPAGE™ 4-12% Bis-Tris Protein gel and LY294002 was purchased from Thermo Fisher. Phosphatase inhibitor cocktail set V was purchased from Calbiochem. Anti-insulin receptor beta rabbit polyclonal antibody (sc-711) and mouse monoclonal antibody (sc-57342) were purchased from SantaCruz. Anti-Akt mouse monoclonal antibody (Cat. No. 2920), anti-phospho-Akt rabbit monoclonal antibody (Cat. No. 4060) and anti-phospho tyrosine rabbit monoclonal antibody (Cat. No. 8954) were purchased from Cell Signaling Tech. Anti-phosphotyrosine mouse monoclonal antibody (PY20) was purchased from BD Transduction Lab. IRDye 800CW donkey antimouse antibody and IRDye 680RD donkey anti-rabbit antibody were purchased from LI-COR biosciences.

### Plasmids

The ACP sequence from the ACP-GPI plasmid [25] was incorporated to the HIR plasmid previously described [24] by the Gibson assembly (New England Biolabs). The HIR (Homo sapiens, NCBI Reference Sequence: NM_001079817) with the *HindIII* site before the start codon and the *XbaI* site after the stop codon, was subcloned by replacing the EGFP sequence of pEGFP-N1 (Clontech, Takara) [24]. The sequences of the HIR, the backbone plasmid, pEGFP-N1 lacking the EGFP sequence, and the ACP were amplified by PCR with PrimeSTAR Max DNA Polymerase (Takara). The primers for PCR were listed in the S1 Table. The KD 2031-ACP-IR plasmid was generated by a point mutation at Lys^1045^ to Arg (AAG to AGG). The fragment sequence of HIR, the *XhoI* site at 3020 to the stop codon, in the plasmid of WT 2031-ACP-IR was replaced to the *XhoI-XbaI* fragment of HIR sequence including the K1045R mutation with the *XbaI* site after the stop codon synthesized by GeneArt (ThermoFisher). The sequences of plasmids were verified and in the case of plasmids generated by Gibson Assembly all sequences including the plasmid backbone were verified by sequencing by Fasteris SA.

### Cell Culture and stable CHO cell lines expressing ACP-tagged human insulin receptor

Parent CHO-K1 and CHO-K1 cells stably expressing extracellular ACP-tagged human insulin receptors were maintained at 37°C with 5% CO_2_ in HAM’s F-12 medium (Thermo Fisher) supplemented 10% fetal bovine serum (FBS), 100 U/ml penicillin and streptomycin. CHO-K1 cells were transfected with ACP-human IR plasmid using Xfect transfection reagent (Takara). 1 day after transfection, the cells were selected in 600 μg mL^-1^ G418, and stably expressing clones were isolated by limiting dilution method. The clones were screened by staining with CoA-Cy3. Cells stably expressing ACP-IR were maintained in the growth medium supplementing with 600 μgmL^-1^ G418.

### Double staining of CoA-PEG11-NR and DAPI

CHO cells stably expressing ACP-IR were washed with serum-free HAM’s F-12 medium and incubated for 30 min at room temperature in serum-free HAM’s F-12 medium with 1 μM CoA-PEG11-NR, 1 μM SFP synthase,10 mM MgCl_2_ and 15 mM HEPES. After washing the excess staining solution with PBS, cells were fixed with 3% paraformaldehyde in PBS, then stained with 300 nM DAPI for 20 min at room temperature. Imaging of the labelled cells was performed under a Zeiss LSM 780 confocal microscope equipped with Plan-Apochromat 63X (1.40 NA) oil immersion objective.

### Emission spectrum measurement of NR12S or CoA-PEG-NR linked to the ACP-IR

For labeling with NR12S, cells stably expressing ACP-IR cultured on glass bottom (Mat Tek) dishes for 2 days were labeled with 800 nM of NR12S for 7 min at room temperature in serum-free medium including 15 mM HEPES after washing with serum-free medium. Stably expressing ACP-IR cells were labelled with 2 μM of CoA-PEG-NRs in serum-free medium containing 15 mM HEPES with 1 μM of SFP synthase and 10 mM MgCl_2_ at room temperature for 20 min. Images were captured using lambda mode under a Zeiss LSM 780 confocal microscope (Zeiss) equipped with plan-apochromat 63 X oil immersion objective (1.4 NA) and 32-channel GaAsP detectors, excitation at 514 nm and detection with 15 channels at 561-695 nm (step: 8.9 nm). All images were 4-time line averaged. Cellular cholesterol-modification was performed before staining. After cells was washed with serum-free medium, 10 mM of methyl beta cyclodextrin was used in serum-free medium at 37°C for 15 min for cholesterol-depletion, and 2.5 mM of cholesterol-water soluble was incubated in serum-free medium at 37°C for 1 hr for cholesterol-loading. The emission spectrum was measured in number of rois from 5 images per each sample.

Meticulous attention to excitation laser power for time-lapse imaging with our system is required to see clear GP change in the local membrane environment surrounding a membrane protein upon stimulation. GP was decreased by exposure to the excitation laser in a laser-power dependent manner (supplementary fig. 3). We performed time-lapse imaging by excitation at 514 nm with 2.5% of Argon laser power and we demonstrated the differential delta GP upon insulin stimulation to show the effect of insulin on GP change. GP decrease caused by excitation light was eliminated by subtraction of the basal GP change in control treatment (Fig. 5 b, c).

### Voltage sensitivity test

Voltage sensitivity of CoA-PEG-NR attached to ACP-IR was measured as described before [25]. Simultaneous optical and electrophysiological recordings were performed at room temperature on a standard patch clamp setup (HEKA, EPC 10Usb) mounted on an inverted Zeiss Axioobserver A1 equipped with an oil-immersion 40x objective (NA 1.3) after cell labeling with NR12S or CoA-PEG-NR. Cells on the microscope stage were perfused with extracellular bath solution (150 mM of NaCl, 5 mM KCl, 1 mM MgCl_2_, 2 mM CaCl_2_, 10 mM HEPES and 5 mM glucose, pH 7.4 adjusted with NaOH, 300 mOsm). Pipettes had resistance of 3-7 mΩ when filled with intracellular solution (140 mM of CsCl, 10 mM of HEPES, 10 mM of HEDTA, pH 7.35 adjusted with CsOH, 290 mOsm). After establishing whole cell configuration, cell capacitance and access resistance were routinely compensated. All experiments were performed in whole-cell patch clamp configuration. Membrane potential was changed from −60 mV to +40 mV and then back to −40 mV. Fluorescence excitation was delivered using a 100 W Mercury short arc lamp (OSRAM) through BP 540-552 nm (RFP) filter or BP360-540 nm (Rhodamine) filters at 28 mW/mm^2^ power densities measured with an optical power meter at the imaging plane. Fluorescence emission was passed through a LP590 nm (RFP) or BP570-640 nm (Rhodamine) filter and recorded by a Zyla 4.2 Plus (Andor) sCMOS Camera operated with NIS-Elements Ar (Nikon). Fluorescence images were analysed in

NIS-Elements Ar and ImageJ by manually defining the regions of interest (ROI) and calculating the mean unweighted mean of pixel values within this region. The cell-free region was used for calculating background fluorescence, which was subtracted from the cell fluorescence. When required, photobleaching was corrected using an exponential decay function. To determine voltage sensitivity, %Δ F/F values were generated by subtracting baseline fluorescence at holding potential from fluorescence during the voltage step and plotted against voltage. Voltage sensitivity of different substrates was then compared as %Δ F/F per 100 mV depolarization from a −60 mV to +40 mV voltage step.

### Western blotting

3.0 × 10^5^ of cells stably expressing WT 2031-ACP-IR or KD 2031-ACP-IR were cultured on 6-well plates for 2 days. After 2 hr serum-starvation, cells were stimulated with 100 nM of human reconstitute insulin for 5 min at 37 °C. The cells were lysed with 120 μl of lysis buffer (50 mM Tris-HCl (pH7.5), 150 mM NaCl, 1 mM EDTA and 0.5% TritonX-100) containing protease- and phosphatase-inhibitor cocktails and incubated on ice for 20 min, then the lysate was centrifugation at 14,000 rpm for 20 min at 4°C. The supernatant was collected and the protein concentration was determined by BCA protein assay. 20 μg of the lysates were heated at 65 °C for 10 min in SDS-sample buffer (6X SDS sample buffer, 375 mM Tris-HCl (pH6.8), 12% SDS, 40% glycerol, 0.03% bromophenol blue and 0.6 M DTT) and subjected to SDS-PAGE on NuPAGE 4-12% Bis-Tris protein gel, then the proteins on the gel were transferred to a nitrocellulose membrane. After overnight blocking of the membrane with 10% BSA in TBST (10 mM Tris-HCl (pH8.0), 150 mM NaCl, 0.05% Tween-20), the membrane was probed with primary antibodies, anti-insulin receptor beta rabbit polyclonal antibody and anti-phospho-tyrosine mouse monoclonal antibody, or anti-insulin receptor beta mouse monoclonal antibody and anti-phospho-tyrosine rabbit polyclonal antibody at 4°C for overnight, followed by secondary antibodies, IRDye 800CW antimouse antibody and IRDye 680RD anti-rabbit antibody, then imaged used Fusion FX (VILBER). For experiments to assess the effect of attachment of PP-PEG11-NR on the receptor tyrosine activity of 2031-ACP-IR, cells were incubated with 1 μM of SFP synthase and 10 mM of MgCl_2_ in 15 mM HEPES containing medium in the presence and absence of 2 μM CoA-PEG11-NR at room temperatures for 30 min after the serum-starvation.

### Time-lapse imaging of the local membrane environment surrounding the insulin receptor

Cells stably expressing WT 2031-ACP-IR cultured on glass bottom dishes for 2 days were labeled with 2 μM of CoA-PEG5-NR, CoA-PEG11-NR, 3μM of CoA-PEG27-NR or 800 nM of NR12S at room temperatures. KD 2031-ACP-IR stably expressing cells plated on glass bottom dishes were labeled with 2 μM of CoA-PEG5-NR. Cells transiently expressing ACP-GPI placed on glass bottom dishes were stained with 2 μM CoA-PEG11-NR. The excess dye was washed with 37°C serum-free HAM’s F-12 medium 3 times, and 1 ml of 37°C serum-free medium was added to the dish, then the dish was placed on the stage top heating chamber (Tokai Hit, INUBTF-WSKM) warmed at 37 °C set on the stage of a Zeiss LSM 780 confocal microscope 5 min before the observation. Spectrum images in the range of 561-695 nm were captured with 15 channels (step 8.9 nm, 4 times line-average) using lambda mode excited at 514 nm every 10 sec for 9 min, and 100 μl of serum-free medium (control) and 100 μl of 1 μM human reconstitute insulin in serum-free medium (final conc. 100 nM) were added to the dish using a syringe with the long needle at 60 sec and 240 sec, respectively, and either of the control or the insulin solution was added at 120 sec for 7 min observation. WT 2031-ACP-IR stably expressing cells labeled with CoA-PEG5-NR were used for inhibitor experiments, and cell staining and observation were performed in the presence of DMSO (control), 0.2 μM of Larunculin B or 50 μM of LY294002.

### Image analysis for time-lapse imaging of the local membrane environment surrounding the insulin receptor

#### Image segmentation of the receptor-linked environment-sensitive probe

Image analysis was performed on all time-lapse data sets to reveal how the GP value changes during insulin signaling. The captured images, however, usually include information such as image noise that is not relevant to the observation targets. Therefore, image segmentation was performed to extract the target regions, the receptor-linked environment-sensitive probe. The detailed procedures are as follows:

1. Image sequences of eight spectral channels from 565 to 645 nm (i.e., 565, 575, 585, 595, 615, 625, 635, and 645 nm) were collected throughout the time-lapse observation.
2. A median filter with a kernel size of w = 3×3×3 was used to filter each time-lapse image sequence.
3. A difference of Gaussian (DoG) filter [40] was performed to each median filtered image.
4. The z-score normalization was performed to the DoG filtered image sequence.
5. K-means clustering [41] was performed to the normalized image sequence in order to form two classes (target probe or background noise).

The DoG filter enhances the edges in a digital image and we performed that filter with a standard deviation of 15 and kernel window size *W* of 25. As the outputs of K-means clustering, we obtained two-class labelled images, showing to which class belonged each pixel. The class with the higher mean value was identified as the target probe class, and only the pixels belonging to this class were used to calculate the GP value.

#### The calculation of GP value, delta GP and differential delta GP

The value of each pixel in the images detected by our method was converted into a GP value according to the following equation [21]:

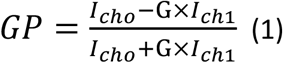

Here, G is the calibration factor, which is calculated with a theoretical GP value of the dye in DMSO measured by a fluorometer and an experimental GP value of the dye in DMSO measured by the same microscope setup and settings as those used for the time-lapse observation [18, 21]. The G factor is defined by the following equation:

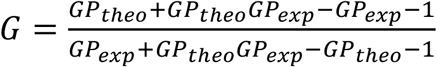

We used GP_theo_= −0.735 and −0.728 for NR12S and CoA-PEG-NR, respectively. GP_exp_ was measured and the G factor was calculated every time when the time-lapse observation was done. Images *I*_ch0_ and *I*_ch1_ of the equation (1) are the averaged images of the lower four channels (565-595 nm) and the higher four channels (615-645 nm), respectively. The translated GP values vary between −1.0 (disordered) and 1.0 (ordered). We depict them as pseudocolor GP images (Fig. 4 a, supplementary fig.5 a) which reveal the reaction of the target probes surrounding the receptor to the insulin solution.

In order to compare time-dependent GP changes, the delta GP at each time was estimated by subtraction of the mean GP at time 0 using R scripts and the mean of the delta GP from multiple time-lapse observations was plotted by Graphpad Prism. For 7 min observation of the KD 2031-ACP-IR and inhibitor treatments, the differential delta GP upon insulin stimulation was calculated by subtraction of the basal delta GP with control treatment from the means of delta GP with insulin treatment at each time point. The basal delta GP was the mean of delta GP from control-treated multiple time-lapse data set and the mean of differential delta GP from multiple time-lapse observations was plotted (Fig.5 b, c).

#### Dynamic analysis of change in GP using zero-mean normalized cross correlation

Although these pseudocolor GP images provide us with information regarding the range of the GP values, the changes in the GP value are quite small throughout the entire time-lapse observation. As a result, we are not able to visualize the point at which the GP values change before and after the application of insulin solution in these images. Moreover, the GP value of each pixel contains the fluctuation caused by measurement noise or the resolution limit of the microscope.

To solve these problems and to visualize the differences in GP values in detail, we utilized a similarity measurement based on zero-mean normalized cross correlation (ZNCC) [42] to reveal the slight changes in GP value in a local region. ZNCC is often used to measure the similarity between two images, and is calculated according to the following equation:

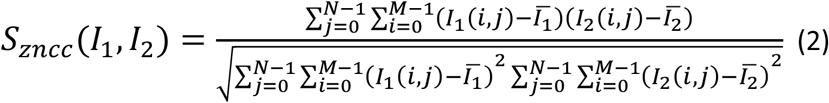

Here, *I*_1_ and *I*_2_ denote the images of a local *M* × *N* region, and 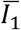 and 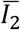 denote their corresponding mean intensities.

ZNCC is known as a measurement that is robust to changes in brightness. Hence, it is able to measure the slight differences in the intensity distribution between the local two regions from a statistical viewpoint. The use of ZNCC in this study enables us to visualize minute differences in the GP values despite the limitations of the imaging. To measure the local similarity of the GP images obtained from Eq. (1), the parameters of ZNCC were set as *M* = 19 and *N* = 19. The time-lapse image sequences were constructed from 55 images (from t = 1 to t = 55). Within those time series, we defined the following reference image for calculating ZNCC. The reference image was created by averaging the part of time-lapse images, from t=7 to t=24, which were the images during the control treatment (Supple fig. 5 b). The differences in the GP values of each pixel at time *t* are calculated as follows:

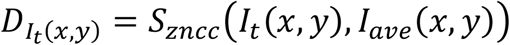

where *I_ave_* is the averaged image used as the reference of ZNCC calculation. The value of D ranges from −1 to 1, where 1 represents an exact match. We did not calculate the differences in the GP value for t≤ 7 because this was before the control solution was applied. The ZNCC values are visualized for the segmented regions (that is, the target probe pixels) as pseudocolor images after median filtering (with a kernel size w = 3 × 3). By analyzing these images, we are able to understand the distinct regions in which the GP changes occurred. According to sort the ZNCC value in each time-lapse image, we found the top 1000 pixels and the bottom 1000 pixels, showing the biggest and the smallest ZNCC changes, respectively (Fig. 4c and supple fig.5 c). As for the representative regions showing the biggest and the smallest ZNCC changes, we created the kymograph images, which shows the dynamic changes of ZNCC during the time-lapse observation (Fig. 4d).

## Supporting information

Movie time lapse GP

Movie time lapse ZNCC

Movie time lapse top bottom

## Acknowledgements

We would like to thank members of the Riezman lab for their help and comments on the manuscript. This work was funded by the NCCR Chemical Biology and the Swiss National Science Foundation (grants 51NF40 -185898, 310030_184949).

