## Supplemental figures and table for "Receptor-linked environment-sensitive probe monitors the local membrane environment surrounding the insulin receptor"

Supplemental information

Figure legends

#### Supplementary figure 1. Comparison of emission spectra of CoA-PEG5-NR linked to 2031-ACP-IR in the time-course

Emission spectrum of a representative time-course observation of CoA-PEG5-NR linked to 2031-ACP-IR (Fig. 3C) at time 0 sec, at 230 sec, before insulin treatment, and at 540 sec, the last time point.

#### Supplementary figure 2. The whole time-lapse image set of Fig. 4 and enlarged image set of ZNCC and Top/Bottom 1000 pixels.

The whole time-lapse image set of GP (a), ZNCC (b) and top/bottom 1000 pixels (c) in Fig.4. Enlarged images of ZNCC time-lapse images (d) and top/bottom 1000 pixels (e).

#### Supplementary video 1-3.

The time-lapse movies of GP (1), ZNCC (2), and top/bottom 1000 pixels (3). Arrow heads point out the regions often forming membrane ruffling or protrusions.

#### Supplementary figure 3. Strong laser power induces GP decrease and masks the effect of insulin

Time course of mean delta GP change compared to time 0. After 2 hr serum starvation, cell stably expressing 2031-ACP-IR or transiently were labelled with CoA-PEG5-NR. Spectral imaging was performed using different laser powers and control solution or insulin solution were treated at 120 sec (upper; weaker laser power) or at 180 sec (lower; stronger laser poser). Data represent the mean ± S.E.M of from 10 time-lapse image sets for both control and insulin of stronger laser powers, 11 for control of lower laser power and 13 for insulin of lower laser power.

#### Supplementary figure 4. Akt phosphorylation is blocked by LY294002 treatment upon insulin stimulation

AKt-phosphorylation upon insulin stimulation in the presence or absence of LY294002. 2031-ACP-IR expressing cells were serum-starved for 2 hr followed by incubation with serum-free medium, medium including 50 μM of LY294002 or DMSO for 10 min at 37 degrees, then cells were stimulated by 100 nM of insulin at 37 degrees for 5 min and phosphorylation of Akt was accessed by western blotting analysis using α-Akt antibody (upper bands) and α phospho-Akt antibody (2[nd] upper bands). Actin was used as a loading control (lower bands).

#### Supplementary figure 5. Actin cytoskeleton disruption by Latrunculin B

Actin cytoskeleton was disrupted by Latrunculin B treatment. Cells plated on glass-bottom dishes were stained with Sir-Actin at 37 degrees for 1 hr, followed by washing excess Sir-Actin, then cells were treated by DMSO (upper) or 0.2 μM of Latrunculin B (lower) at 37 degrees for 10 min. Left pictures are Nomarski images and right pictures are images of Sir-Actin.

#### Supplementary figure 6. Emission spectrum of ACP-GPI

a) Emission spectrum shows the local membrane environment of 2031-ACP-IR is more disordered than that of ACP-GPI. Cells transiently expressing 2031-ACP-IR were stained with CoA-PEG11-NR or NR12S and emission spectrum between 561 and 695 nm was measured from 10 images for CoA-PEG11-NR (n=119 regions of interest (rois)) and from 5 images for NR12S (n=75). Cells transiently expressing wild type ACP-GPI were stained with CoA-PEG11-NR and emission spectrum between 561 and 695 nm was measured from 5 images (n=75). Normalized fluorescent intensities were plotted. b) Our system recognizes the difference in the local membrane environment between wild type ACP-GPI and PGAP2/3 mutant ACP-GPI. Cells transiently expressing wild type ACP-GPI or PGAG2/3 mutant ACP-GPI were stained with CoA-PEG11-NR and emission spectrum between 561 and 695 nm was measured from 6 images (n=35 for wild type and n=36 for PGAP2/3 mutant). c) The emission spectrum of the bulk plasma membrane assessed by NR12S was identical between wild-type ACP GPI expressing cells and PGAP2/3 mutant ACP-GPI expressing cells. The data represents mean ± S.E.M from 6 images (n=36 rois for each sample).

### Methods

#### Materials

Anti-Akt mouse monoclonal antibody (Cat. No. 2920) and anti-phospho-Akt rabbit monoclonal antibody (Cat. No. 4060) were purchased from Cell Signaling Tech. Sir-Actin was purchased from Spirochrome and verapamil was purchased from Sigma-Aldrich.

#### Sir-Actin staining in living cells

CHO cells stably expressing ACP-2031-IR were incubated with HAM's F-12 HAM medium containing 3 μM of Sir-Actin and 10 μM of verapamil at 37 degrees for 1 hr, followed by washing with HAM's F-12 medium 3 times. Imaging of the labelled cells was performed using a Leica SP5 confocal microscope equipped with HCX PL APO CS 63 X oil immersion objective (1.4 NA).

### Supplementary figure 1

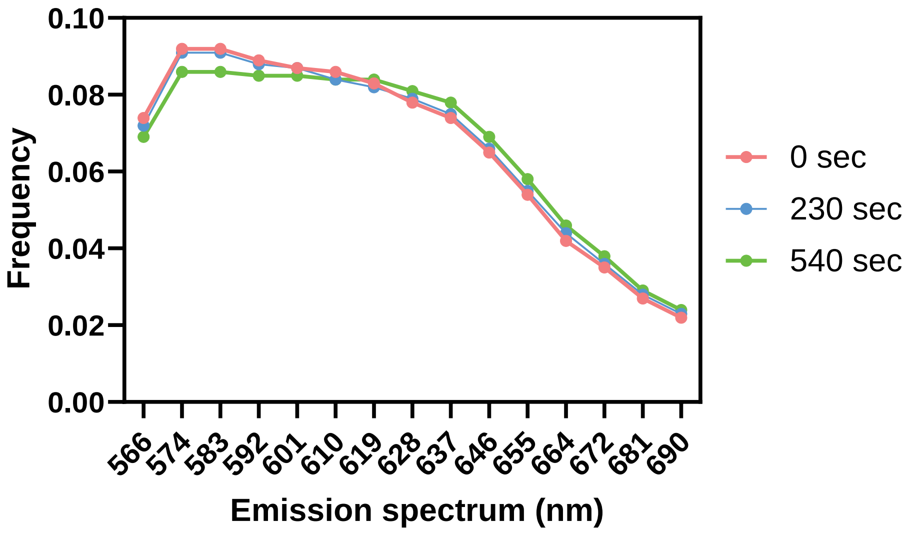

### Supplementary figure2 a

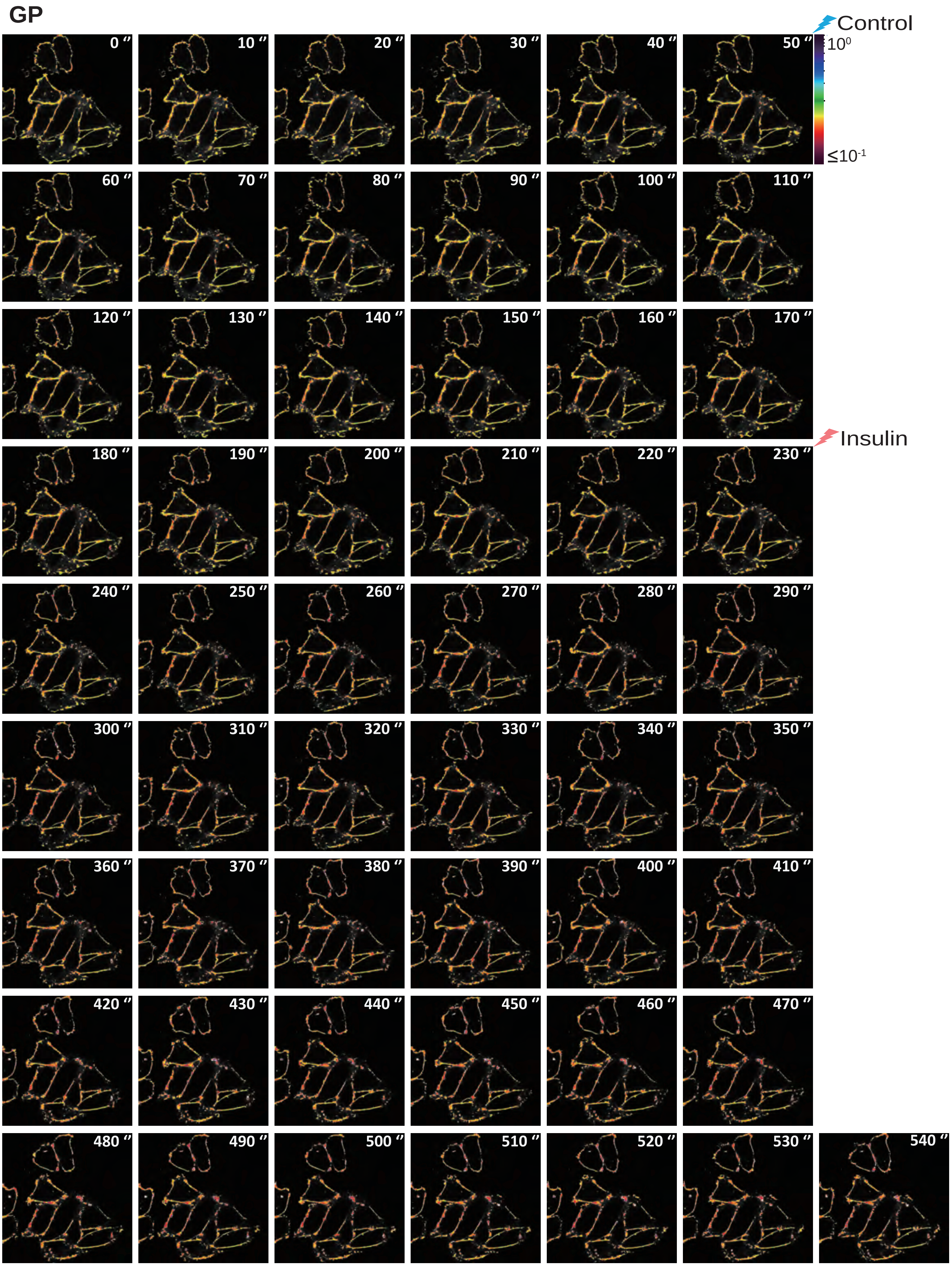

### Supplementary figure 2 b

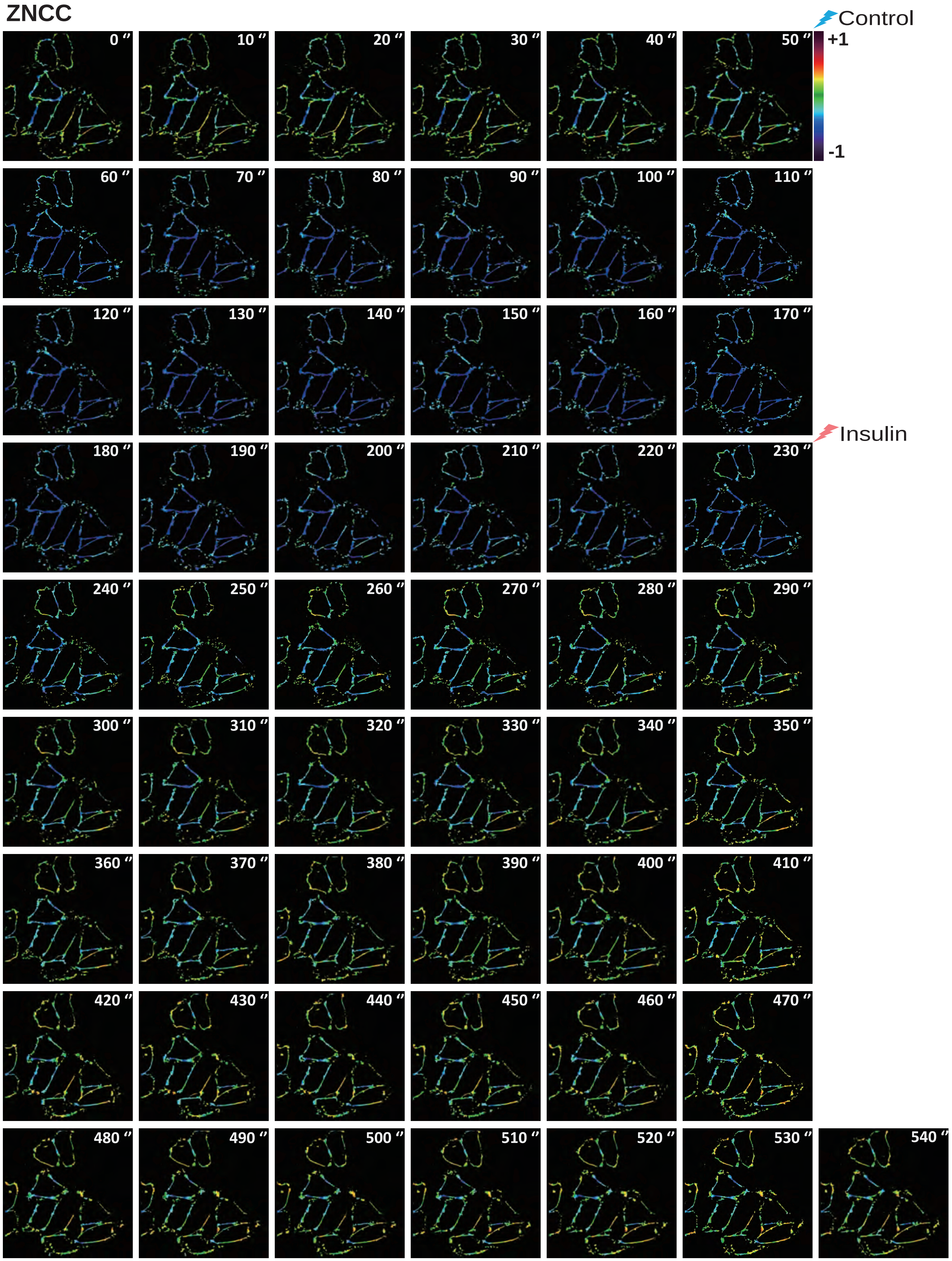

### Supplementary figure 2 c

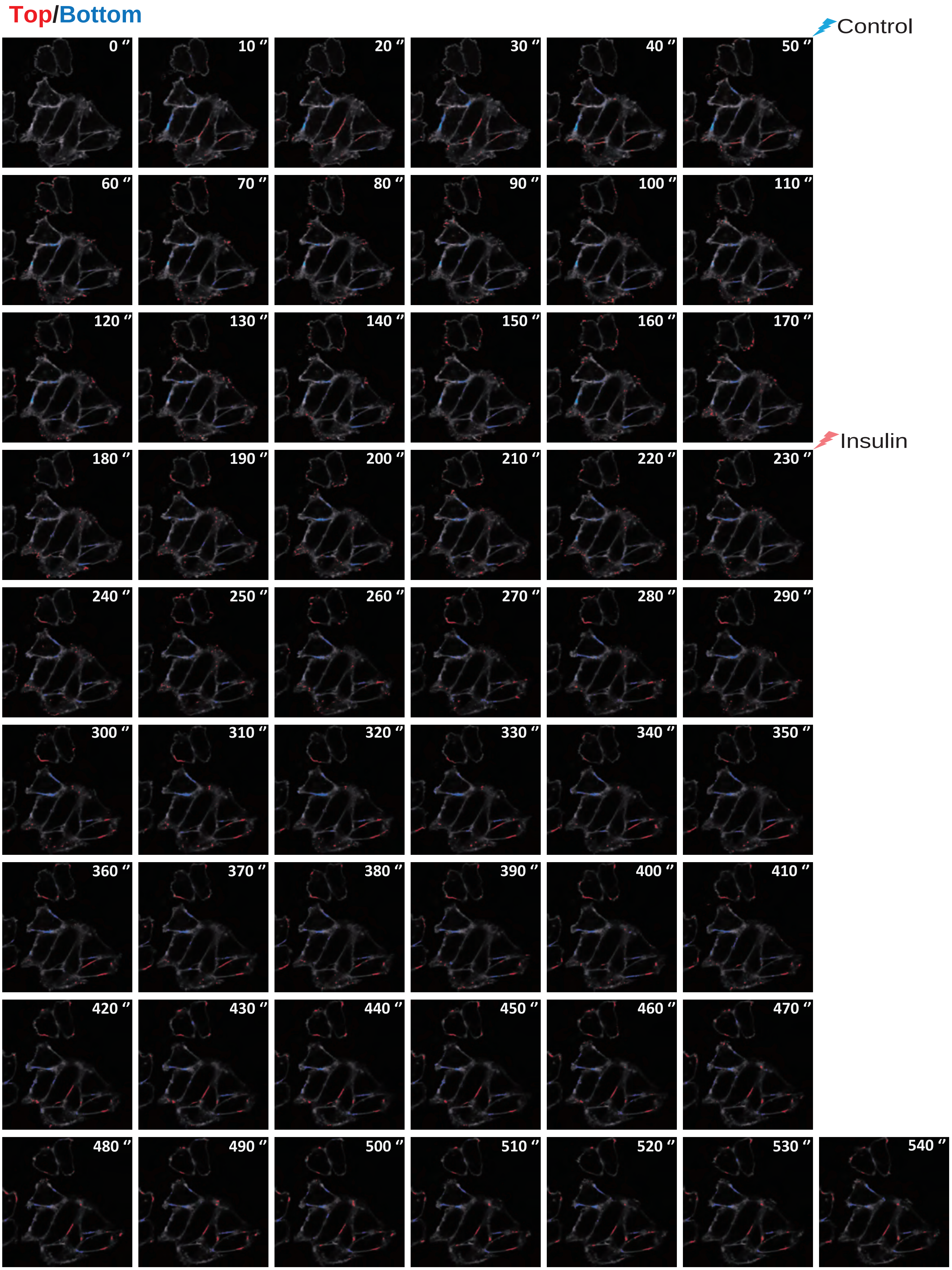

### Supplementary figure 2d

#### ZNCC

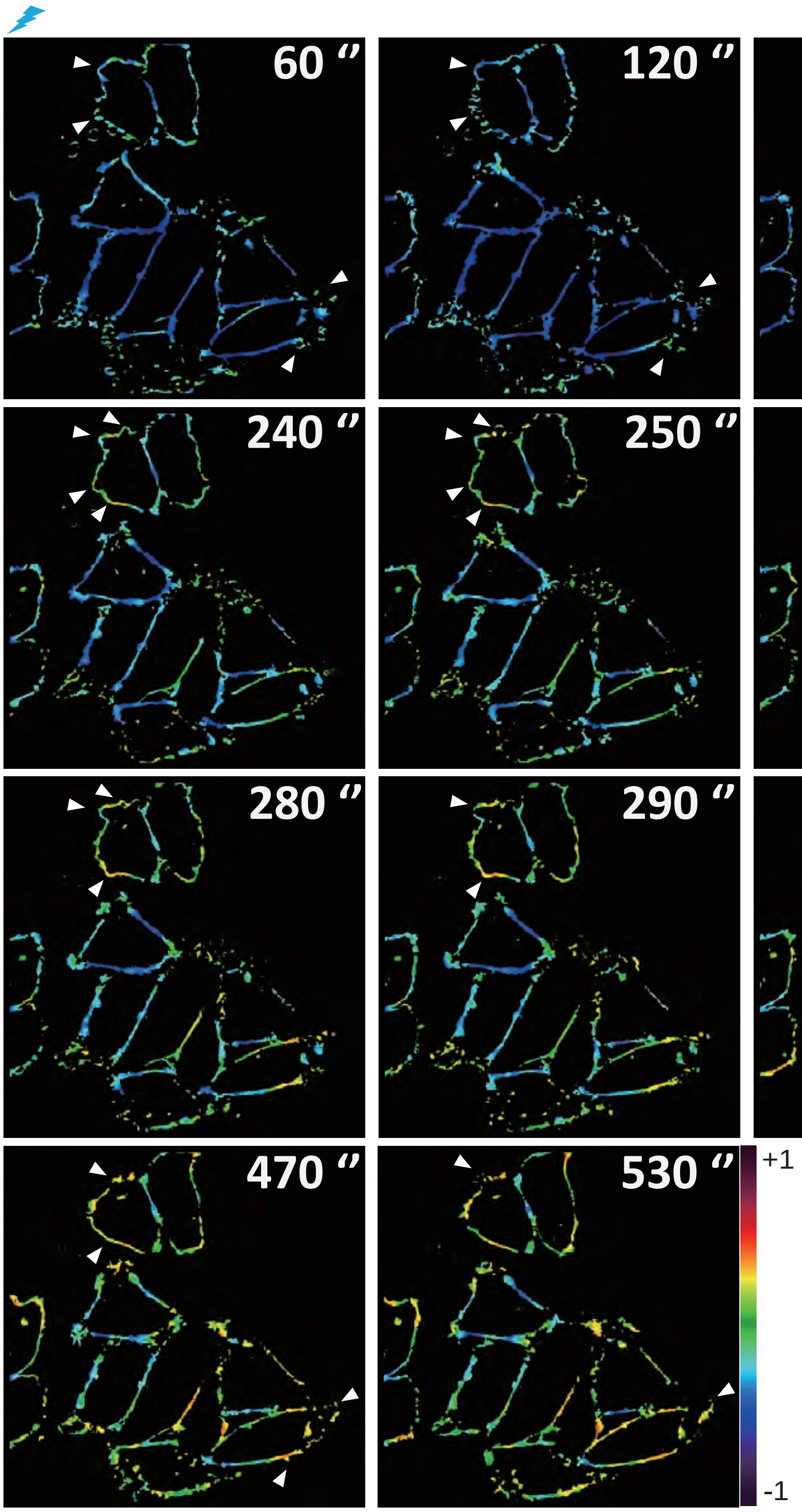

#### Supplementary figure 2e

Top/Bottom

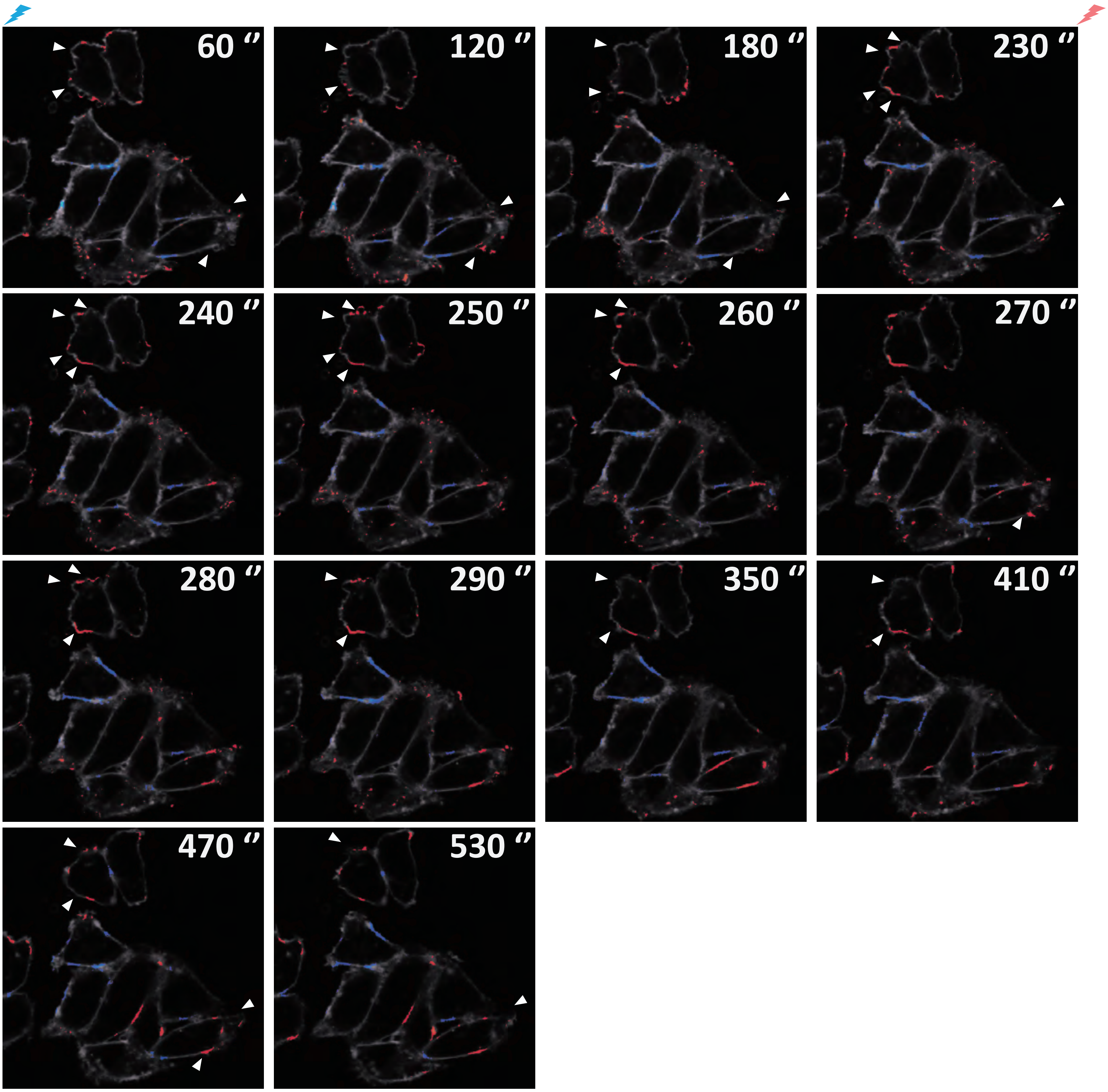

### Supplementary figure 3

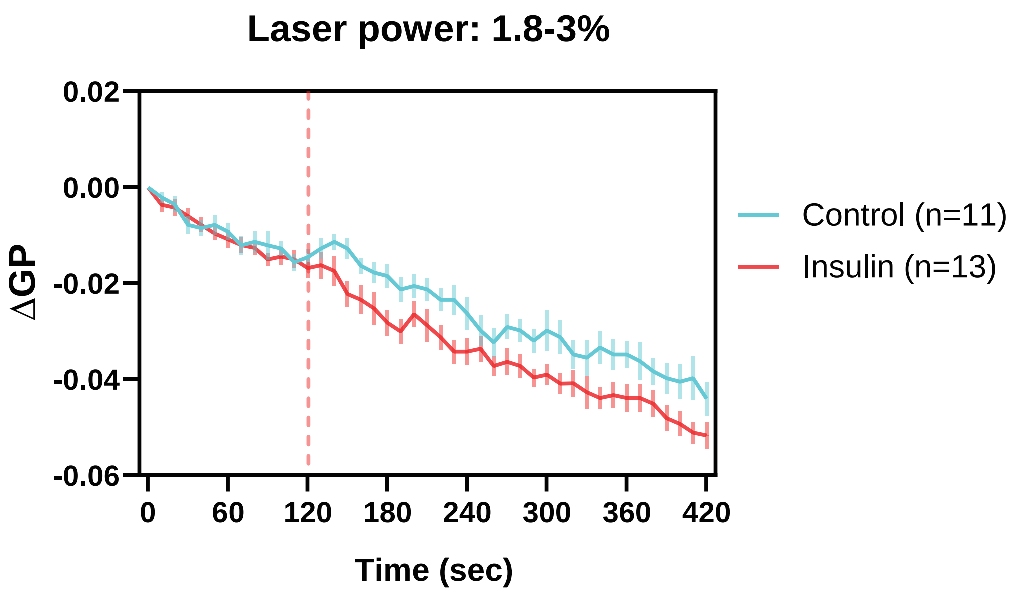


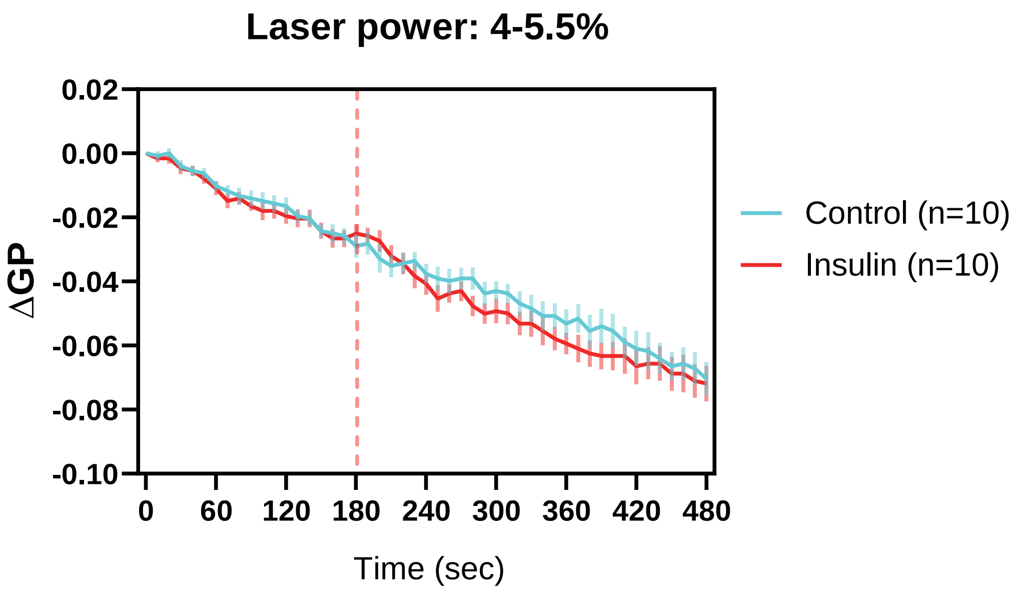

#### Supplementary figure 4

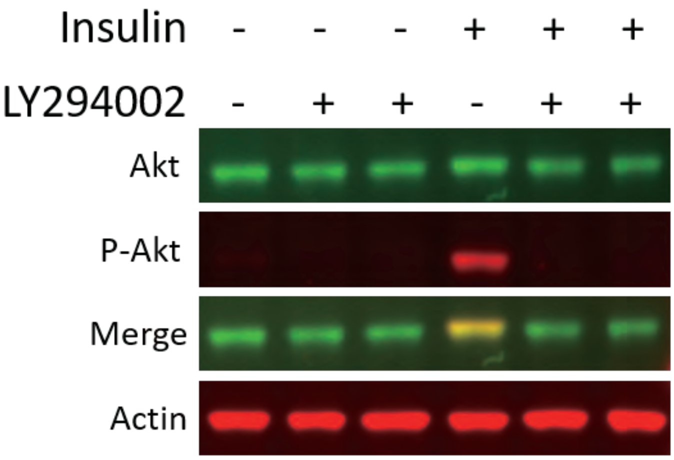

#### Supplementary figure 5

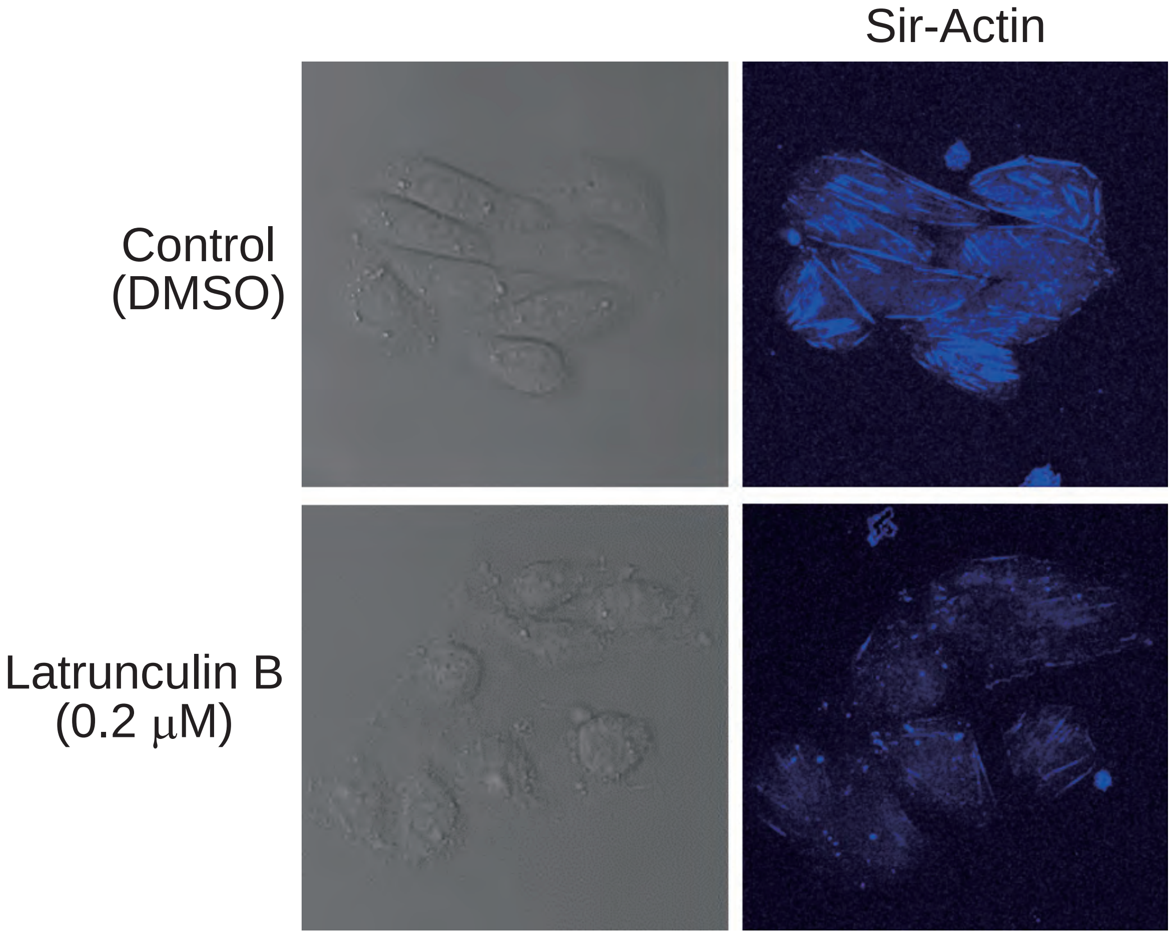

**Supplementary figure 6**

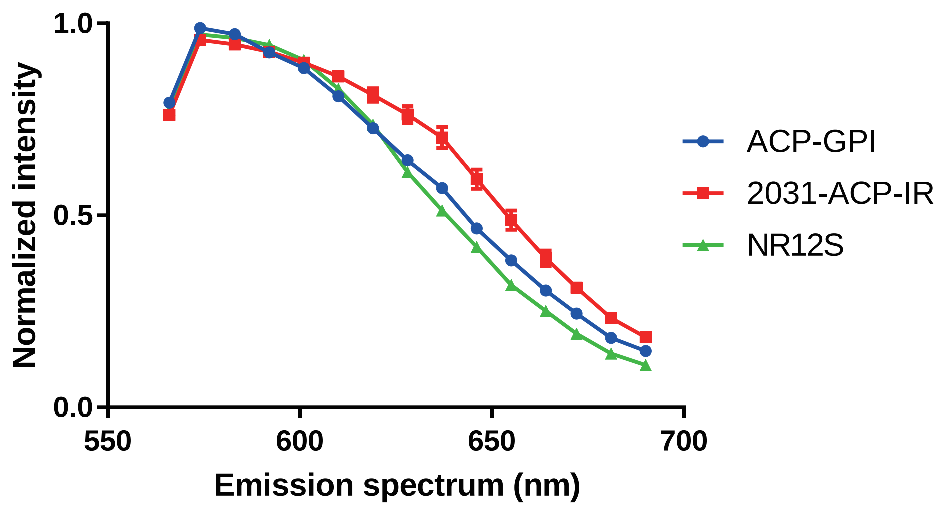


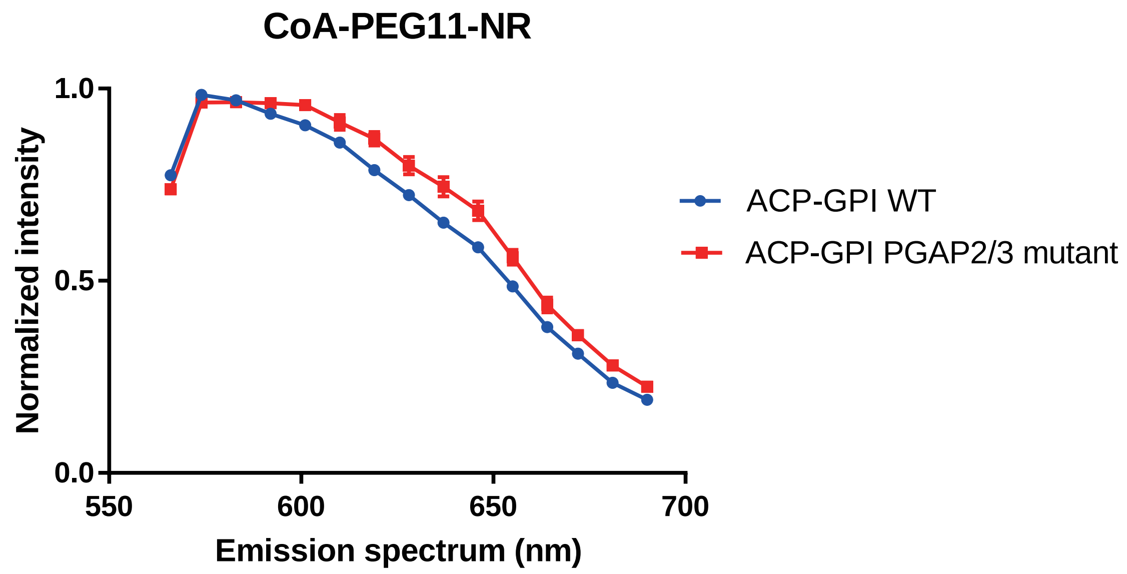


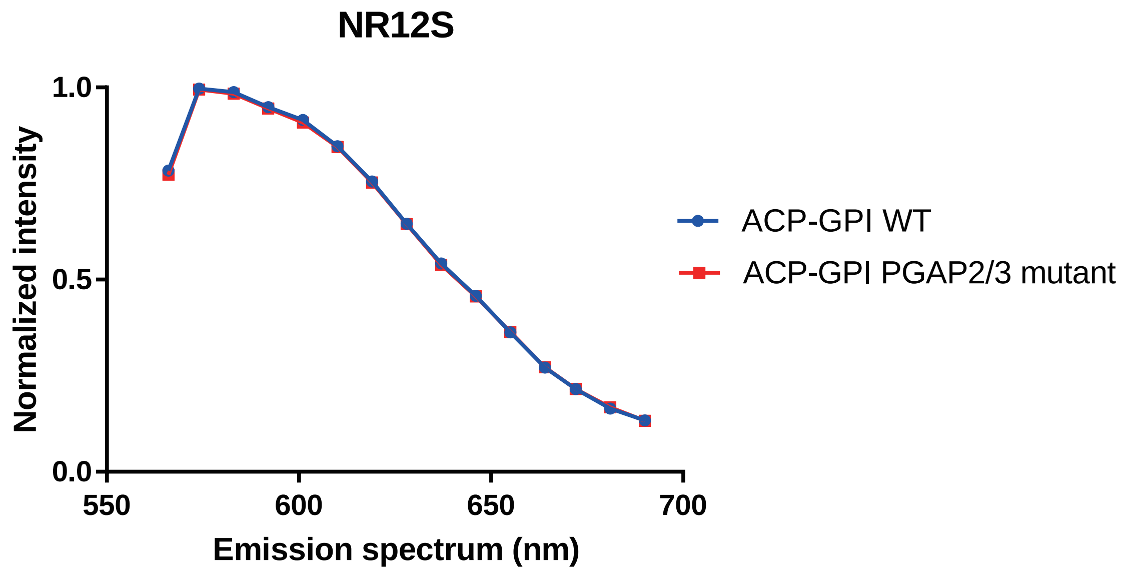

### Supplementary Information

**Supplementary Table 1 |** Summary of the primers used in this study.

| Plasmid name | Primer | Sequence | Template |
|---|---|---|---|
| Plasmid backbone | pEGFPN1_Gibon F | TCT AGA TCA TAA TCA GCC ATA CCA CAT TTG TAG | pEGFP-N1 |
| Plasmid backbone | pEGFPN1_Gibon R | AAG CTT GAG CTC GAG ATC TGA G | pEGFP-N1 |
| 1992-ACP-IR | IR_preACP_F | GAC TCA GAT CTC GAG CTC AAG CTT CCG CCA CCA TGG CCA CCG G | Insulin receptor |
| 1992-ACP-IR | 1992-IR_preACP_R | CGA TAG TGC TCA TTT CCG CCT GCC TCT CCC AG | Insulin receptor |
| 1992-ACP-IR | 1992-ACP_F | GAG GCA GGC GGA AAT GAG CAC TAT CGA AGA ACG CGT TAA G | ACP |
| 1992-ACP-IR | 1992-ACP_R | ACA GCT CAC TGT CCG CCT GGT GGC CGT TGA TG | ACP |
| 1992-ACP-IR | 1992-IR_postACP_F | CGG CCA CCA GGC GGA CAG TGA GCT GTT CGA GC | Insulin receptor |
| 1992-ACP-IR | IR_post_R | GTG GTA TGG CTG ATT ATG ATC TAG ATT AGG AAG GAT TGG ACC GAG G | Insulin receptor |
| 2031-ACP-IR | IR_preACP_F | GAC TCA GAT CTC GAG CTC AAG CTT CCG CCA CCA TGG CCA CCG G | Insulin receptor |
| 2031-ACP-IR | 2031-IR_preACP_R | CGA TAG TGC TCA TCC CTT TGA GGC AAT AAT CCA GCT CGA ACA G | Insulin receptor |
| 2031-ACP-IR | 2031-ACP_F | TTG CCT CAA AGG GAT GAG CAC TAT CGA AGA ACG CGT TAA G | ACP |
| 2031-ACP-IR | 2031-ACP_R | AGG GCA GCT TCA GCG CCT GGT GGC CGT TGA TG | ACP |
| 2031-ACP-IR | 2031-IR_postACP_F | CGG CCA CCA GGC GCT GAA GCT GCC CTC GAG G | Insulin receptor |
| 2031-ACP-IR | IR_post_R | GTG GTA TGG CTG ATT ATG ATC TAG ATT AGG AAG GAT TGG ACC GAG G | Insulin receptor |

| | | | |
|---|---|---|---|
| PreCT-ACP-IR | IR_preACP_F | GAC TCA GAT CTC GAG CTC AAG CTT CCG CCA CCA TGG CCA CCG G | Insulin receptor |
| PreCT -ACP-IR | PreCT -IR_preACP_R | CGA TAG TGC TCA TCT TCC TAA ACG AGG ACT CCT CCA GCT C | Insulin receptor |
| PreCT -ACP-IR | PreCT -ACP_F | CTC GTT TAG GAA GAT GAG CAC TAT CGA AGA ACG CGT TAA G | ACP |
| PreCT -ACP-IR | PreCT -ACP_R | AAT CCT CAA ACG TCG CCT GGT GGC CGT TGA TG | ACP |
| PreCT -ACP-IR | PreCT -IR_postACP_F | CGG CCA CCA GGC GAC GTT TGA GGA TTA CCT GCA C | Insulin receptor |
| PreCT -ACP-IR | IR_post_R | GTG GTA TGG CTG ATT ATG ATC TAG ATT AGG AAG GAT TGG ACC GAG G | Insulin receptor |
